# Role of two modules controlling the interaction between SKAP1 and SRC kinases Comparison with SKAP2 architecture and consequences for evolution

**DOI:** 10.1101/2023.12.11.571115

**Authors:** Laurine Levillayer, Camille Brighelli, Caroline Demeret, Anavaj Sakuntabhai, Jean-François Bureau

## Abstract

SRC kinase associated phosphoprotein 1 (SKAP1), an adaptor for protein assembly, plays an important role in the immune system such as stabilizing immune synapses. Understanding how these functions are controlled at the level of the protein-protein interactions is necessary to describe these processes and to develop therapeutics. Here, we dissected the SKAP1 modular organization to recognize SRC kinases and compared it to that of its paralog SRC kinase associated phosphoprotein 2 (SKAP2). Different conserved motifs common to either both proteins or specific to SKAP2 were found using this comparison. Two modules harboring different binding properties between SKAP1 and SKAP2 were identified: one composed of two conserved motifs located in the second interdomain interacting at least with the SH2 domain of SRC kinases and a second one composed of the DIM domain modulated by the SH3 domain and the activation of SRC kinases. This work suggests a convergent evolution of the binding properties of some SRC kinases interacting specifically with either SKAP1 or SKAP2.

## Introduction

SRC kinase associated phosphoprotein family is composed of two members, SKAP1 and SKAP2. Due to their localization to human chromosomes 7p15.2 for SKAP2 and 17q21.32 for SKAP1 at one extremity to respectively HoxA and HoxB cluster, their duplication occurs during evolution of chordates to vertebrates [1]. At least since gnathostome radiation, SKAP1 and SKAP2 are highly conserved. SKAP1 is mainly expressed in the immune system particularly in T-cells [2, 3] in contrast to SKAP2 that is widely expressed [4] even if at higher level in some cells of the immune system such as macrophages and neutrophils [5–8]. SKAP2 also plays an important role during the development of the Central Nervous System [9, 10]. The structure of both proteins is composed of a dimerization domain (DIM) followed by a first interdomain (Link1), a Pleckstrin homology domain (PH), a second interdomain (Link2), and a SRC homology 3 domain (SH3) (Figure 1). The homologous or heterologous association of two DIM domains by hydrophobic interactions formed a four-helix bundle structure that has been solved for SKAP2 [11, 12]. Their PH domain interacts with membranes via two phosphate-derivatives of phosphatidyl inositol, PI(3,4,5)P_3_ and either PI(4,5)P_2_ for SKAP1 [13] or PI(3,4)P_2_ for SKAP2 [12]. Interestingly, the DIM and the PH domains of the same chain interact together with an important role of D120 and D129, respectively for SKAP1 and SKAP2. The interdomain Link2 contains two conserved tyrosine phosphorylated motifs that have been suspected to bind to the SH2 domain of SRC kinases [14]. Both SH3 domains bind proline-rich motif of PRAM1 [15], FYB1 [4, 16], FYB2 and FAM102A [17, 18], in which W333 and W336 play a necessary role for SKAP1 and SKAP2, respectively.

**Figure 1:**
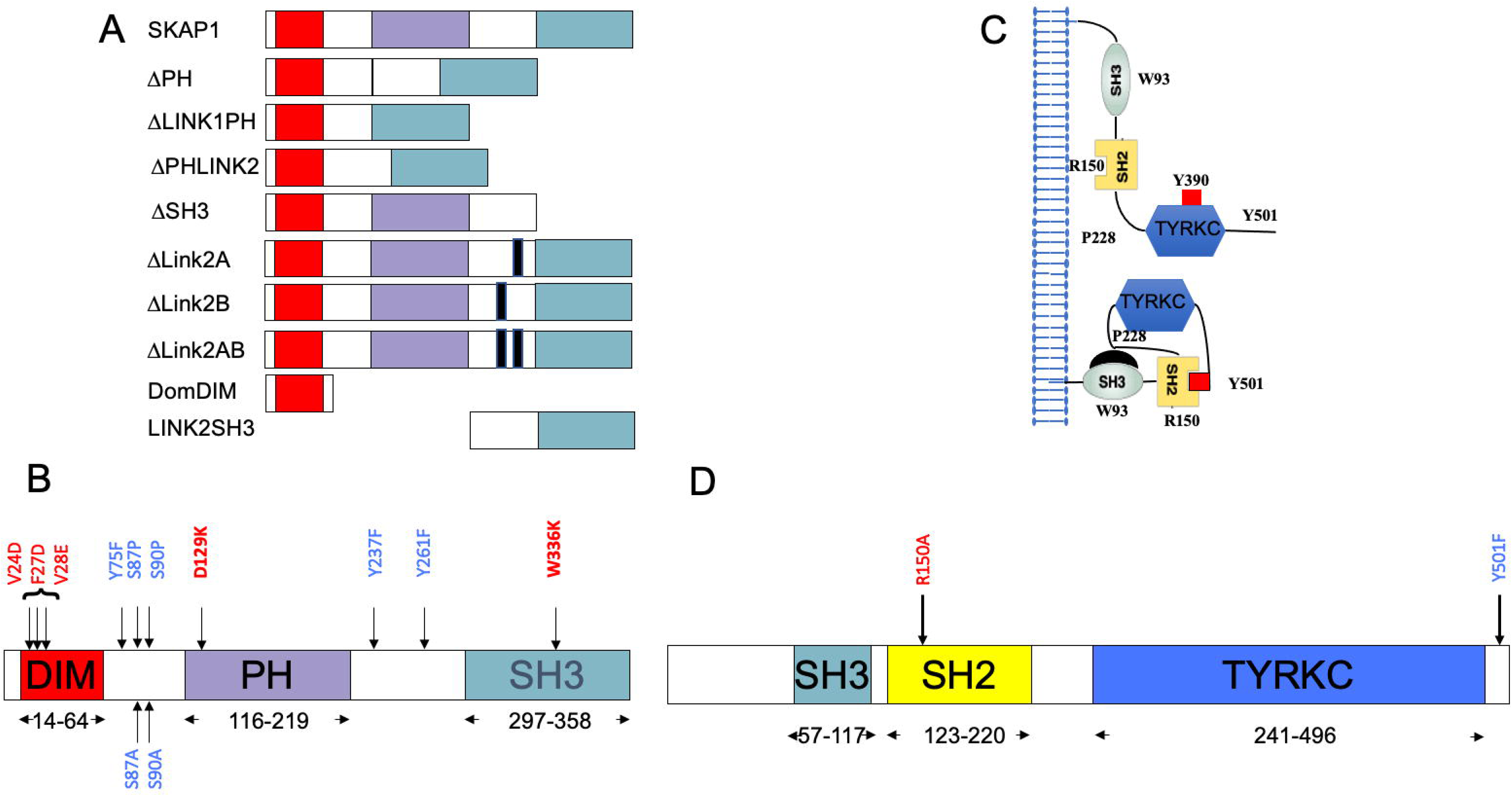
map of SKAP1, SKAP2, and HCK mutants. For each protein, the delineation of domain boundaries is indicated. Wild type and replacement amino acid identities at mutated position involved in domain or motif inactivation are indicated in red and blue respectively. (A) Schematic of SKAP1 organization with the three domains, domain of dimerization (DIM), Pleckstrin homology domain (PH), SRC homology 3 domain (SH3), and the two interdomains, Link1 and Link2 with deletion mutants. (B) Schematic of SKAP2 organization with the three domains, DIM, PH, and SH3. (C) Schematic illustrating inactive (bottom) and active (top) forms of HCK. (D) Schematic of HCK protein organization with its three domains: SRC homology 3 domain (SH3), SRC homology 2 domain (SH2) and tyrosine kinase catalytic domain (TYRKC).

At the T-cell immune synapse during the inside-out signaling, SKAP1 stabilizes SLP-76/LCP2 microclusters by interacting with FYB1 [19, 20]. Interestingly, the interaction of either SKAP1 or SKAP2 with FYB1 protects them from degradation [21]. SKAP1 also formed a complex with LAT and GRB2 induced by the integrin LFA-1 and PTK2A/PTK2B, which terminate T-cell conjugate formation [20]. During this process, SKAP1 is recruited to lipid raft, in which the SRC kinase FYN phosphorylate its tyrosine 271, a necessary event for its interaction with GRB2 [14]. Note also that Skap1 interacts by its DIM domain with Rapl and RIAM, regulators of respectively α and β-integrin [19, 22, 23]. SKAP1 was also reported to interact with Polo-like kinase 1 for an optimal control of cell-cycling [24]. A recent review [25] discuss these data in more details. The FYB1-SKAP1 complex has also been detected in mast cells but its function is unknown [26]. In conclusion, both SKAP1 and SKAP2 play an important role as adaptors for the same proteins such as FYB1, FAM102A, and SRC kinases. Our recent work revealed that the binding of SRC kinases by SKAP2 is dictated by its complex modular architecture, inviting to study the SKAP1 one [27].

The SRC kinase family is composed of the SrcA subfamily with FGR, FYN, SRC, and YES, the SrcB subfamily with BLK, HCK, LCK, and LYN and finally the FRK-related family with FRK, SRMS, and PTK6 [28, 29]. SRC kinases [30], which phosphorylate tyrosine residues, are composed of three domains, a SRC homology 3 (SH3) domain, a SRC homology 2 (SH2) domain, and a tyrosine kinase catalytic domain (TYRKC) (Figure 1D). A fine allosteric control of the kinase activity is modulated from the passage of a closed to open conformation [31–33](Figure 1C). The closed, inactive conformation is provided by intramolecular interactions between (1) the SH3 domain and a proline-rich motif located in the hinge between the SH2 and the catalytic kinase domains and (2) the SH2 domain and a phosphorylated tyrosine motif in the C-terminus of the kinase. In the open active conformation, these intramolecular interactions are lost, and a transphosphorylation of a tyrosine located in the active loop of the catalytic site locks the system [34]. The localization of most SRC kinases in different membranous compartments is finely regulated by N-terminal myristylation and palmitoylation [35, 36]. However, neither SRMS nor FRK have an N-terminal myristylation site.

Evolution of protein-protein interactions has been a subject of rich debates on the gain and loss of interactors particularly after a duplication event [37–42]. Alternative splicing also plays an important role to explain tissue-specificity of an interaction and of monogenic disorders [43–45]. Also, evolution information was frequently used to predict interaction [38]. Large protein and domain families have been particularly studied [46]. Kinases with an SRC module constitute one family in which such analysis was performed [47–49]. This kind of analysis was also conducted on domains such as the SH2 and the SH3 domains [50–53]. One interesting case was protein-protein interactions between viral proteins and host ones [54]. Most of the works were done at the global level of the interactome but some of them also analyzed the level of protein-protein interfaces [38, 55, 56]. Knowledge at this level is necessary to develop protein-protein inhibitors [57, 58]. Specific cellular expression occurs among some SRC kinases as for SKAP1 and SKAP2 allowing to test possible convergent evolution. The FYN kinase is particularly interesting because one isoform, FYNT, is mainly expressed in T lymphocytes as SKAP1, and a second one, FYNB, is expressed in the brain as SKAP2 [59]. Both isoforms differ by their exon 7 sequence, which were absent to the FYN isoform that we used in our previous work [60, 61]. Interestingly, mis-expression of these two isoforms in T lymphocytes as in the brain has been associated to an increasing risk for some diseases [62]. LCK kinase is also mainly expressed in T lymphocytes in contrast to HCK kinase, which is expressed in the monocytes/macrophages lineage and neutrophils as SKAP2.

In the present paper, we described SKAP1 architecture recognizing SRC kinases, and compared it to that of its paralog SKAP2 [27]. Comparison of similar deletion mutants and mutations between these two adaptors allowed to precisely define conserved and divergent functions. We identified a motif in Link1 interdomain, specific to SKAP2, which affects the interactions with SRC kinases. The conservation of the interdomain Link2 function between both proteins allowed the detection of two common conserved motifs, which play non-redundant and possibly cooperative roles in the binding to the SH2 domain of SRC kinases. These two conserved motifs and at least the SH2 domain of SRC kinases constitute the first interaction unit identified; the second one is composed of the DIM domain, modulated by its SH3 domain and the activated SRC kinases. These two modules in SKAP1 and SKAP2 harbor different binding properties to SRC kinases. Interestingly, LCK and FYNT kinases expressed in T lymphocytes as SKAP1 show different binding properties than HCK kinase expressed in the monocytes/macrophages lineage and neutrophils, and FYNB kinase expressed mainly in the brain. SKAP2 is also mainly expressed in the monocytes/macrophages lineage, in neutrophils and in the brain suggesting a possible coevolution between these two adaptors and SRC kinases.

## Materials and Methods

### Plasmids, mutagenesis, BP and LR cloning, and luciferase complementation assay

Previously used plasmids and a more detailed methodology will be found elsewhere [17, 27]. The ORF of *FYNT* (NM153047.4), *FYNB* (NM002037.5) flanked by two gateway sites have been synthesized and cloned in PUC57 plasmid (GenScript, Hong Kong). These two clones were individually transferred by Gateway recombinational cloning into pDONR207 vector. The ORF of the new SKAP2 deletion mutants flanked by two gateway sites were amplified by PCR, cloned into pCR-II TOPO vector using TOPO TA cloning kit (Thermo Fisher Scientific, Waltham) and transferred into pDONR207 vector. Mutagenesis of SKAP1, SKAP2, and HCK ORFs were performed using QuickChange Lightening Site-Directed Mutagenesis kit (Agilent Technologies, Les Ulis) according to the manufacturer’s protocol. Supplementary Table S1 shows the new primers used for all these purposes. The sequence of other primers will be found in previous publications [17, 27]. After each mutagenesis and PCR amplification, the insert was completely sequenced at Eurofin services using LightRun Barcodes (Eurofins Genomics, Ebersberg, Germany). Sequence alignments were performed using DNA Strider [63]. The ORF of SKAP1, and SKAP2 mutants were transferred into pSPICA-N2 vector and those of SRC kinases and HCK mutants into pSPICA-N1 and pSPICA-C1 ones. The pSPICA vectors are mammalian expressing vectors designed for luciferase complementation assay [64]. They expressed *Gaussia princeps* Luciferase fragment 1 (amino acid residues from 18 to 109) or 2 (amino acid residues from 110 to 185) at the N or C-terminal part of the fusion protein. The luciferase complementation assay was performed as described in human HEK293T cells [17, 64]. Briefly the first day, 30 000 HEK293T cells are cultured in 100 μl of DMEM supplemented with 10% Fetal Bovine Serum and antibiotics per well of microplate 96 wells (Greiner, Kremsmünster). One day later, cells were transfected with 100 ng GPCA plasmid pair using polyethylenimine (PEI) method. The third day, culture medium was discarded, cells were washed once with 150 μl of Phosphate Buffer Saline without calcium and magnesium (PBS) and incubated for 25 min in 40 μl of lysis buffer. Luminescence monitoring was performed after addition of native Coelenterazine on a Centro XS^3^ LB 960 microplate luminometer (Berthold Technologies, Thoiry). Transfections using PEI were performed at least in triplicate. Protein-protein interactions were monitored by measuring interaction-mediated normalized luminescence ratio (NLR) [64]. Briefly, NLR is the ratio between the interaction signal and a control one, the sum of signals of each component of the interaction with the complementary empty hemi-luciferase expressing vector.

### Experimental Design and Statistical Rationale

Approximation of the mean and the variance of a ratio were performed using Taylor expansions with a null covariance between the numerator and the denominator. Two-sample z test and two-way ANOVA after log-normal transformation were used to compare these ratios. Differential interaction scatterplots and PPI-mutation plots were performed as described in [17]. Three levels of significance were used for the P-value lower than 0.05 (*), 0.01 (**), and 0.001 (***). Bonferroni correction was used for multiple testing with an initial threshold p = 0.05. To compare different experiments, we used modified NLR for each SRC kinases as the ratio between NLR of the SKAP1 mutant and NLR of wild-type SKAP1 protein, which was named wild-type normalized NLR. Experiments were repeated three times. For studying the effect of the different domains and mutations, we used the ratio between NLR of the SKAP1 mutant and NLR of the same SKAP2 mutant, which was named SKAP2 normalized NLR. Study of each mutant experiment was repeated at least two times. Statistical analyses were performed using Stata/IC version 10.1 (Stata Corporation, College Station, USA). Comparison of mean to theorical value (i.e. Mean ratio to 1 and its log-normal transformation to 0) and between two means was performed by using Student t test and unpaired Student t test, respectively. Comparison of mean to theorical value was also studied using binomial distribution after calculating the number of samples upper/lower to theoretical value. Comparison of two means was also performed by Kruskal-Wallis test. Parametric and non-parametric tests giving similar results, only results of the parametric are shown in Table1. We also decided to present preferentially ratio data that are easier to interpret than those after log-normal transformation when both analyses gave similar results. The only exception was the two-way ANOVA, which was performed after a log-normal transformation of the modified NLR and using analytic weights. For data modelling, the full model contains six explanatory variables and the data on the N1- and C1-fused SRC kinases were analyzed separately. Six different explicative variables were generated from the initial experiments on the different mutants. An explicative model was generated from of this full model by a step-by-step procedure that removed at each step the lowest non-significant variable until no non-significant variables were present. A principal componant analysis (PCA) from the present mutant data and those of SKAP2 was performed after a log-transformation using the R package Factominer [65] and Factoshiny according to the author specification to determine which factors explained the variability amongst C1-fused SRC kinases. More details on the methodology, complete results, and interpretation are shown in supplemental data.

### Conservation of the Casein kinase 1 motif in the interdomain Link1 of SKAP2

The N-terminal fragment of human SKAP2 protein structure was predicted from I-TASSER server [66] using two crystal structures of a N-terminal fragment of murine SKAP2 containing both the helical dimerization domain and the PH domain (PDB ID: **<u>2OTX</u>**and **<u>1U5E</u>**). Clustalw program was used to align respectively 38 protein sequences of SKAP2 from the Coelacanth to mammals. ConSurf server [67, 68] was used to visualize evolutionary conservation in this macromolecule from its predict structure and aligned sequences using a maximum likelihood procedure. UCSF Chimera version 1.13.1 [69] was used to visualize the conservation of these structures using a script from ConSurf server.

## Results

In this paper, we study the SKAP1 organization, which recognizes the SRC kinases. The goal of this work was 1) to define the SKAP1 modular architecture of this interaction; 2) to specify the composition and binding properties of its two modules by looking for common or specific motifs to SKAP1 and SKAP2 architecture; 3) to compare the two architectures in the context of evolution. For that purpose, we used a luciferase complementation assay in HEK293T cells where the C-terminal hemi-gaussia luciferase (Gluc2) is fused at the N terminus of SKAP1 (N2-SKAP1) and the N-terminal hemi-gaussia luciferase (Gluc1) is fused to either the N terminus (N1) or C terminus (C1) of the SRC kinases. The two orientations of hemi-gaussian luciferase fused to SRC kinases were studied because we previously showed that they affect their subcellular localization and activation but also the interactions with SKAP2 [27]. By similarity of what we found for SKAP2 adaptor [27], our assumption was that there is a general SKAP1 architecture, which defines its interaction with all the SRC kinases. We measured each PPI signal by using the normalized luciferase ratio [64] and the mean value between SRC kinases of each mutant to detect the general architecture. In practice, to assess the effect of SKAP1 deletions or mutations, the NLR generated by their binding to an SRC kinase has been divided by the binding of wildtype SKAP1 to the same kinase. This generated a wild-type normalized NLR enabling to standardize the interaction of mutated SKAP1 for each SRC kinase amongst the different experiments. The SKAP1 interaction with nine SRC kinases, BLK, FGR, FRK, FYN, HCK, LCK, LYN, SRMS, and YES, was studied with 12 SKAP1 mutants, nine of which are deletion mutants (Fig. 1A) and the others contain nonsynonymous mutations. To compare architecture between SKAP1 and SKAP2, we also studied similar SKAP2 mutations for their interaction with the nine SRC kinases. Figure 1B shows the localization of all SKAP2 nonsynonymous mutations. Lastly, we studied the specifity of some SRC kinase interactions with SKAP1 and SKAP2 by the analysis of two nonsynonymous mutations on HCK (Fig. 1D) or LCK background and of the three FYN isoforms, and more systematically by using a Principal Component Analysis (PCA) on C1-fused SRC kinases and either SKAP1 or SKAP2 mutant data.

### SKAP1 modular architecture interacting with SRC kinases

We first studied the interaction of nine SKAP1 mutants with nine SRC kinases, which has been able to previously define the SKAP2 architecture [27]. Figure 2 shows one experiment for each SKAP1 mutant. The interaction of SRC kinases systematically decreases for ΔPHLink2, DomDIM1, and Link2SH3 SKAP1 mutants, independently of the localization of the hemi-gaussia fusion. This strongly suggests that the domain DIM and the interdomain Link2 are necessary and sufficient for the interaction with the SRC kinases. The Link2SH3 and the DomDIM mutants have a similar loss of signal indicating that neither the interdomain Link2 nor the DIM domain are sufficient. The ΔPH, the ΔLink1PH, and the ΔSH3 mutants do not exhibit such decrease of the PPI signal supporting that they are not necessary. The W333K SKAP1 mutant, invalidated for the binding to the proline-rich binding motif, behave similarly as the ΔSH3 mutant, showing a slight increase in their binding capacity. The D120K and DIM^-^ mutations have a slight effect on the PPI respectively increasing the signal for C1-fused SRC kinases and decreasing it for N1-fused ones.

**Figure 2:**
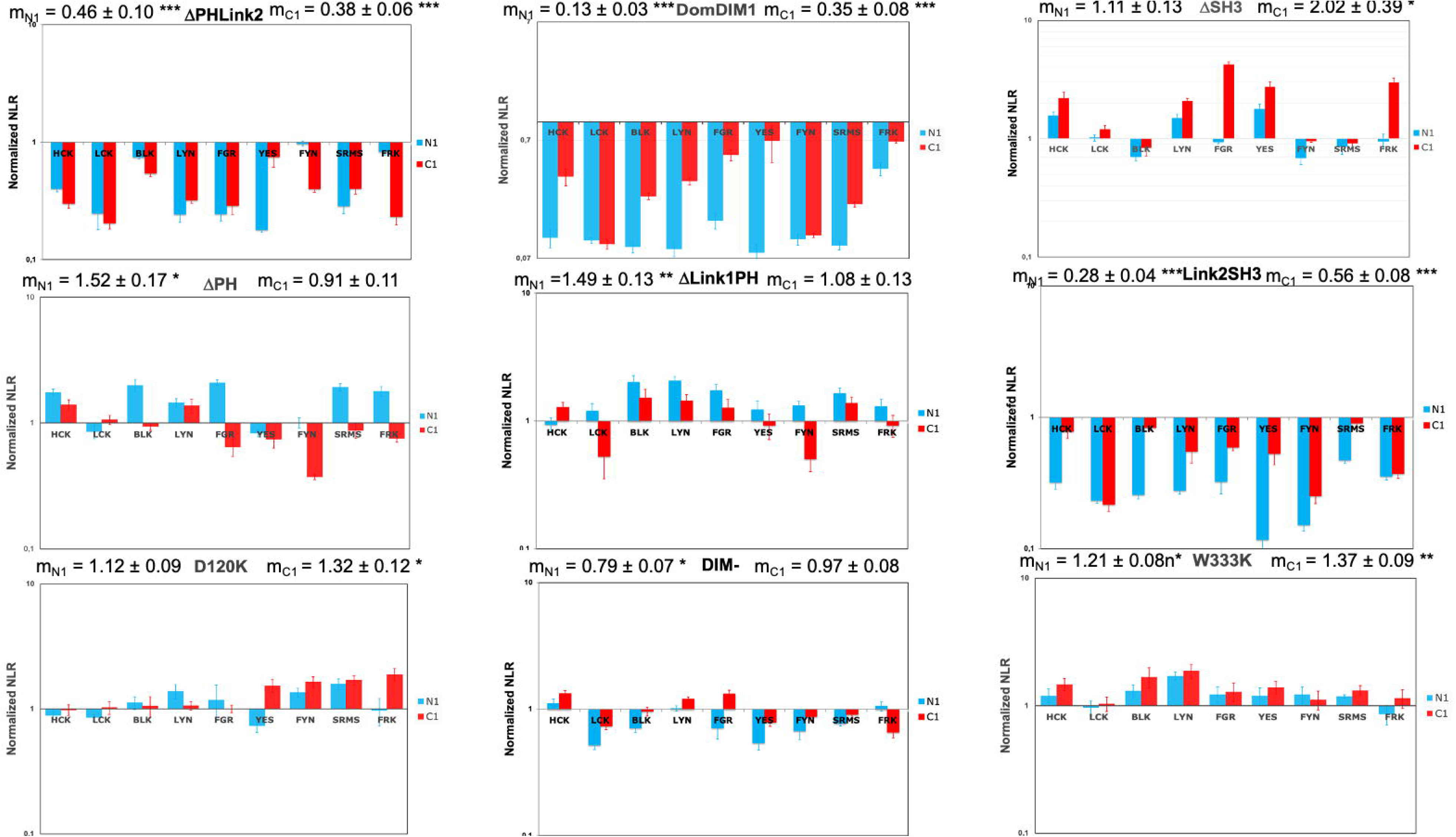
Study of the interaction between SKAP1 mutants and N1-fused or C1-fused SRC kinases. Graphs show wild type normalized NLR of nine SKAP1 mutants with N1-fused (blue) and C1-fused (red) SRC kinases in a logarithmic scale. Wild type normalized NLR is the ratio of the NLR between a SKAP1 mutant and one SRC kinase to the corresponding NLR between wild type SKAP1 and the same SRC kinase fused with hemi-gaussia at the same position. For each graph, the normalized NRL mean ± SEM among SRC kinases is shown as well as its level of significance (*p* = 0.05 (*), *p* = 0.01 (**), *p* = 0.001 (***)).

In conclusion, our data suggest a SKAP1 binding architecture independent of its SH3 domain and of the status of the SRC kinases (i.e the localization of the hemi-luciferase), which clearly differs from the one we identified for SKAP2 using the same strategy [27].

### Statistical validation of the model of SKAP1 architecture

Previous hypotheses were confirmed using univariate analysis combining three experiments. For each SKAP1 mutant, three mean-value effects were tested on N1-fused SRC kinases and on C1-fused ones comparing to theorical mean of no effect equal to one and on the difference between the signal of the C1-fused SRC kinases and that of N1-fused ones. Results were considered statistically significant only after Bonferroni’s correction (Table 1).

**Table 1:**
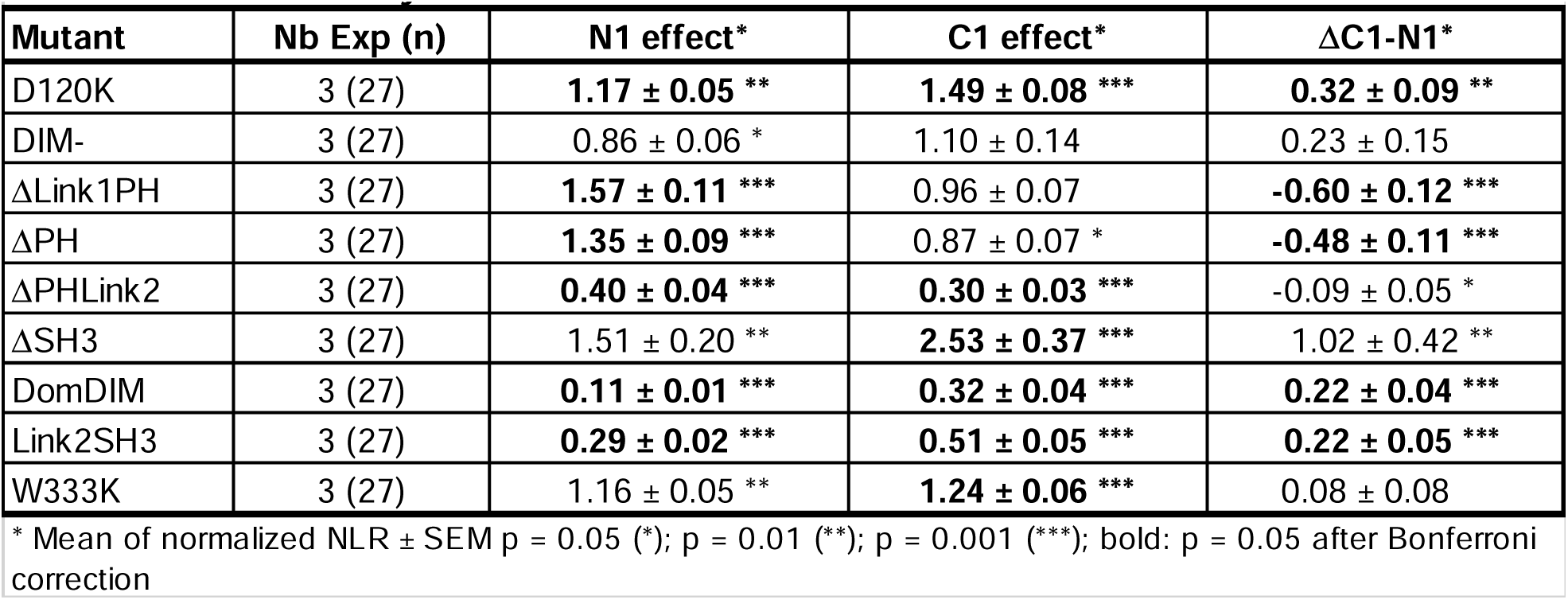
Statistical analysis of the interaction between SKAP1 mutants and SRC kinases.

| Mutant | Nb Exp (n) | N1 effect* | C1 effect* | ΔC1-N1* |
| --- | --- | --- | --- | --- |
| D120K | 3 (27) | **1.17 ± 0.05 \*\*** | **1.49 ± 0.08 \*\*\*** | **0.32 ± 0.09 \*\*** |
| DIM- | 3 (27) | 0.86 ± 0.06 * | 1.10 ± 0.14 | 0.23 ± 0.15 |
| ΔLink1PH | 3 (27) | **1.57 ± 0.11 \*\*\*** | 0.96 ± 0.07 | **-0.60 ± 0.12 \*\*\*** |
| ΔPH | 3 (27) | **1.35 ± 0.09 \*\*\*** | 0.87 ± 0.07 * | **-0.48 ± 0.11 \*\*\*** |
| ΔPHLink2 | 3 (27) | **0.40 ± 0.04 \*\*\*** | **0.30 ± 0.03 \*\*\*** | -0.09 ± 0.05 * |
| ΔSH3 | 3 (27) | 1.51 ± 0.20 ** | **2.53 ± 0.37 \*\*\*** | 1.02 ± 0.42 ** |
| DomDIM | 3 (27) | **0.11 ± 0.01 \*\*\*** | **0.32 ± 0.04 \*\*\*** | **0.22 ± 0.04 \*\*\*** |
| Link2SH3 | 3 (27) | **0.29 ± 0.02 \*\*\*** | **0.51 ± 0.05 \*\*\*** | **0.22 ± 0.05 \*\*\*** |
| W333K | 3 (27) | 1.16 ± 0.05 ** | **1.24 ± 0.06 \*\*\*** | 0.08 ± 0.08 |

\* Mean of normalized NLR ± SEM p = 0.05 (\*); p = 0.01 (\*\*); p = 0.001 (\*\*\*); bold: p = 0.05 after Bonferroni correction

Comparison of the deletion mutants shows that (1) only three mutants, ΔPHLink2, DomDIM, and Link2SH3, have a significant decrease of the PPI signal; (2) other mutants except DIM^-^ one have at least one significant increase of the PPI signal; (3) two mutants, ΔLink1PH and ΔPH have a significant increase of the N1-fused signal with a significant interaction; (4) two mutants, W333K and ΔSH3, have a significant increase of the C1-fused signal; (5) D120K mutant has a significant increase of N1-fused and C1-fused signal significantly greater for C1-fused one. These results validate the model of SKAP1 architecture in which the domain DIM and the interdomain Link2 are necessary and sufficient for binding to SRC kinases independently of the location of hemi-gaussia fusion. In addition, the similar interaction pattern of ΔLink1PH and ΔPH mutants suggests that the PH domain decreases probably indirectly the interaction with N1-fused SRC kinases. The increase of binding to SRC kinases for D120K mutant supports either a role of the intramolecular interaction between the DIM and the PH domains [12] or the appearance of a new binding site for lipids by increasing the interaction of the PH domain to membranes [12, 13].

A statistical modeling was performed taking account both literature curation and our results. Six variables (see Table 2) were generated, describing function possibly affected by mutations and a status for each variable was assigned for each mutant. However due to overfitting, these six variables cannot be analyzed together, and we decided to suppress the variable DIM^-^. Note that the loss of PH domain combined ΔLink1PH and ΔPH mutant data. A two-way ANOVA analysis was carried out after lognormal transformation of the dependent variable on these five variables with either N1-fused or C1-fused SRC kinase data. This transformation was chosen since most data are linearized after log-log transformation in differential interaction scatterplot (see supplemental Fig. S1). An explicative model was generated from a full model by a step-by-step procedure that removed at each step the lowest nonsignificant variable until no nonsignificant variable was present (Table 2).

**Table 2:** Statistical modeling of the interaction between SKAP1 and SRC kinases.

|  |  |  |  |  |  |  |  |  |  |  |
| --- | --- | --- | --- | --- | --- | --- | --- | --- | --- | --- |
| Variable | Partial SS | df | N1-Loc |  |  | Partial SS | df | C1-Loc |  |  |
|  |  |  | MS | F (sense) | P |  |  | MS | F (sense) | p* |
| Explanatory model | 146.81 | 5 | 29.36 | 130.05 | $7.20 \cdot 10^{-66}$ | 142.21 | 4 | 35.55 | 136.10 | $2.62 \cdot 10^{-60}$ |
| Loss of PH domain | 5.26 | 1 | 5.26 | 23.75 (+) | $2.01 \cdot 10^{-6}$ | | | | | NS |
| D120K | 2.08 | 1 | 2.08 | 9.21 (+) | 0.0027 | 4.44 | 1 | 4.44 | 16.99 (+) | 0.0001 |
| ΔSH3 | 1.41 | 1 | 1.41 | 6.23 (+) | 0.0132 | 33.19 | 1 | 33.19 | 127.07(+) | $6.74 \cdot 10^{-24}$ |
| W333K | 1.61 | 1 | 1.61 | 7.14 (+) | 0.0081 | 1.35 | 1 | 1.35 | 5.17 (+) | 0.0239 |
| SKAP1 module | 28.46 | 1 | 28.46 | 126.03 | $9.89 \cdot 10^{-24}$ | 37.02 | 1 | 37.02 | 141.72 | $6.04 \cdot 10^{-26}$ |
| Residual | 53.51 | 237 | 0.24 |  |  | 62.17 | 238 | 0.26 |  |  |
| * NS non-significant (deleted variable during the procedure of minimization) |  |  |  |  |  |  |  |  |  |  |

For C1-fused SRC kinase data, four variables were significant, the SKAP1 module consisting of the DIM and Link2 regions necessary and sufficient to bind to SRC kinases, D129K, ΔSH3, and W333K. For N1-fused SRC kinase data, the five variables, the SKAP1 module, D129K, ΔSH3, W333K, and the loss of PH domain were significant. This ANOVA thus corroborates the SKAP1 interaction module as well as the effects of the D120K, W333K, and ΔSH3 mutants as defined in the univariate analysis and supports that the SRC kinase status affects only slightly their binding capacity to SKAP1.

### Comparing SKAP1 and SKAP2 architecture

To analyze the difference between the two architectures more finely, we compared the effect of each mutant and wild type form between SKAP1 and SKAP2. In practice, the NLR generated by the binding of a SKAP1 mutant or wild type SKAP1 to an SRC kinase has been divided by the NLR of same SKAP2 mutant or wild type SKAP2 to the same kinase. This ratio was named SKAP2 normalized NLR. Figure 3 shows the result of one experiment for each of the ten pairwise proteins. For each of the ten data, the mean-value of NLR for the nine SRC kinases was compared to the theorical value of no effect (equal to 1) depending on the localization of the hemi-gaussia fusion. First, we compared the pattern of significant differences between each mutant and the wildtype protein to classify the data. As previously described [17], SKAP1 has a significant decrease of the PPI signal for C1-fused SRC kinases compared to SKAP2. Five mutants, D120K, DIM^-^, ΔPH, ΔLink2PH, and W333K, have a similar pattern suggesting that these deletions or mutations are not responsible for the difference between SKAP1 and SKAP2. Note for the DIM^-^ mutant that its mean of NLR for C1-fused SRC kinase is greater than that for wildtype SKAP1 (m_C1_ = 0.75 versus 0.39). For the ΔSH3 and ΔLink1PH mutants, SKAP1 PPI signals to N1-fused SRC kinases are stronger than the SKAP2 ones. Link2SH3 and DomDIM mutants have a slightly different pattern with no significant difference of signal for N1-fused and C1-fused SRC kinase. From the differential interaction scatterplot between SKAP1 versus SKAP2, the position of each barycenter and the corresponding regression line were calculated (supplemental Fig. S2). The geometric mean of the NLRs from the SKAP1 and SKAP2 data defines the coordinates of the barycenter along the y-axis and x-axis, respectively (Supplemental Fig. S3). If the position of this barycenter varies on a line parallel to the no-effect line with a slope equal to 1, the difference between the NLR mean-values of SKAP1 and SKAP2 will remain constant. All other changes in barycenter position will affect this difference and support that the corresponding mutant has different interactions with SRC kinases than its wild type protein. The position of both barycenters are similar to that of wildtype protein for D120K, ΔPH and W333K mutants. This position of both baricenters moves mainly according to the line of no effect for ΔPHLink2 mutant and probably also for DomDIM one (see C1-fused data and below) suggesting no difference between SKAP1 and SKAP2 data. The position of barycenters for both N1-fused and C1-fused data moves mainly according to the x-axis supporting an effect of ΔLink1PH and ΔSH3 mutants. Only for C1-fused data, a similar x-axis movement is detected for DIM^-^ and Link2SH3 mutants even if we could not exclude a similar effect for DomDIM mutant. In conclusion of these two analyses, ΔLink1PH, ΔSH3, and Link2SH3 mutants show clear differences between SKAP1 and SKAP2 data. Conflictual results have been obtained for DomDIM and DIM^-^ mutants.

**Figure 3:**
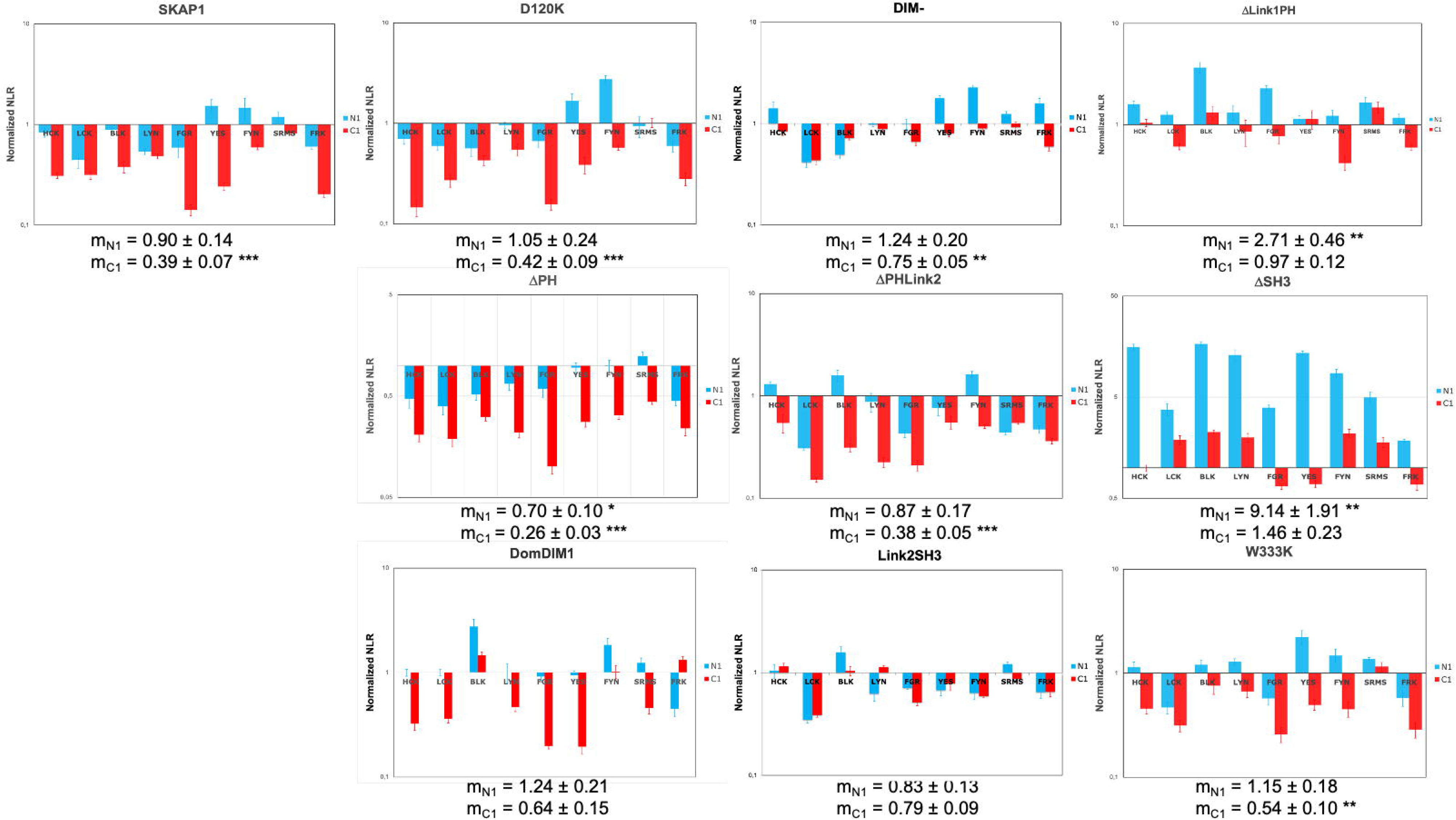
Comparison of the interaction with SRC kinases between SKAP1 and SKAP2 mutants. Graphs show SKAP2 normalized NLR of wild type SKAP1 and nine SKAP1 mutants with N1-fused (blue) and C1-fused (red) SRC kinases in a logarithmic scale. SKAP2 normalized NLR is the ratio of the NLR between a SKAP1 mutant and one SRC kinase to the corresponding NLR between the same SKAP2 mutant and the same SRC kinase fused with hemi-gaussia at the same position. For each graph, the normalized NRL mean ± SEM among SRC kinases is shown as its level of significance (*p* = 0.05 (*), *p* = 0.01 (**), *p* = 0.001 (***)).

A statistical validation was performed by comparing the mean-value of SKAP2 normalized NLR for each mutant with that of wildtype protein for N1-fused or C1-fused SRC kinases. Table 3 shows the results. It confirms that ΔLink1PH and ΔSH3 mutants have a significantly greater mean-value of SKAP2 normalized NLR than that for wildtype protein independently of the localization of the hemi-gaussa fusion. Link2SH3 and DIM^-^ mutants have a significantly greater mean-value of SKAP2 normalized NLR than that for wildtype protein only for C1 data. These results showed that these four mutations have different roles between SKAP1 and SKAP2 since SKAP1 mutant and wildtype SKAP1 NLRs are normalized according the corresponding NLR of SKAP2 mutant and wildtype SKAP2.

**Table 3:** Statistical analysis comparing effect of each mutation between SKAP1 and SKAP2.

| Mutant | Nb Exp (n) | ΔN1-Mutant-N1-SKAP1* | ΔC1-Mutant-C1-SKAP1* |
| --- | --- | --- | --- |
| D120K | 2 (18) | $-0.05 \pm 0.17$ | $-0.01 \pm 0.07$ |
| DIM<sup>-</sup> | 2 (18) | $0.38 \pm 0.18$ | **$0.36 \pm 0.06$** |
| ΔLink1PH | 2 (18) | **$2.00 \pm 0.34$** | **$0.57 \pm 0.10$** |
| ΔPH | 1 (9) | $-0.24 \pm 0.15$ | $-0.10 \pm 0.06$ |
| ΔPHLink2 | 2 (18) | $-0.22 \pm 0.14$ | $-0.04 \pm 0.06$ |
| ΔSH3 | 2 (18) | **$7.39 \pm 1.48$** | **$1.07 \pm 0.22$** |
| DomDIM | 2 (18) | $0.37 \pm 0.18$ | $0.23 \pm 0.16$ |
| Link2SH3 | 2 (18) | $0.03 \pm 0.14$ | **$0.60 \pm 0.10$** |
| W333K | 2 (18) | $0.24 \pm 0.17$ | $0.16 \pm 0.07$ |

\* Difference of means of normalized NLR ± SEM. Bold : $p = 0.05$ after Bonferroni correction

In conclusion, these results strongly suggest that the SH3 domain and the interdomain Link1 have different functions between SKAP1 and SKAP2 proteins. These results support also that dimerization affects differently the interaction to C1-fused SRC kinases between SKAP1 and SKAP2 proteins. Furthermore, the DIM domain has also a major role to explain the differences detected between SKAP1 and SKAP2 proteins for their binding capacity to SRC kinases depending on the localization of the hemi-gaussia fusion. This result explains why the Link2SH3 mutant, which lost the DIM domain, has a significantly greater PPI signal for C1-fused SRC kinases.

### Role of a SKAP2-specific motif located in the interdomain Link1

Our previous results strongly suggest that the interdomain Link1 affects specifically the interaction with SRC kinases for SKAP2. Search for a motif specific to SKAP2 and conserved in its phylogeny identified a motif that has been defined as a site of phosphorylation for Casein kinase 1 [70]. Interestingly, this site is in a α−helix (Fig. 4A), which is absent in SKAP1 such specific to SKAP2. The two serine residues of this site were mutated to either proline or alanine to destroy or not the α−helix. Figure 4B shows that both mutants increased the PPI signal of SRC kinases independently of the localization of hemi-gaussia fusion. This result supports the view that the phosphorylation of this serine affects the interaction with SRC kinases and not the secondary structure. This motif is probably at least one of the elements, which explains the specific role of the interdomain Link1 of SKAP2.

**Figure 4:**
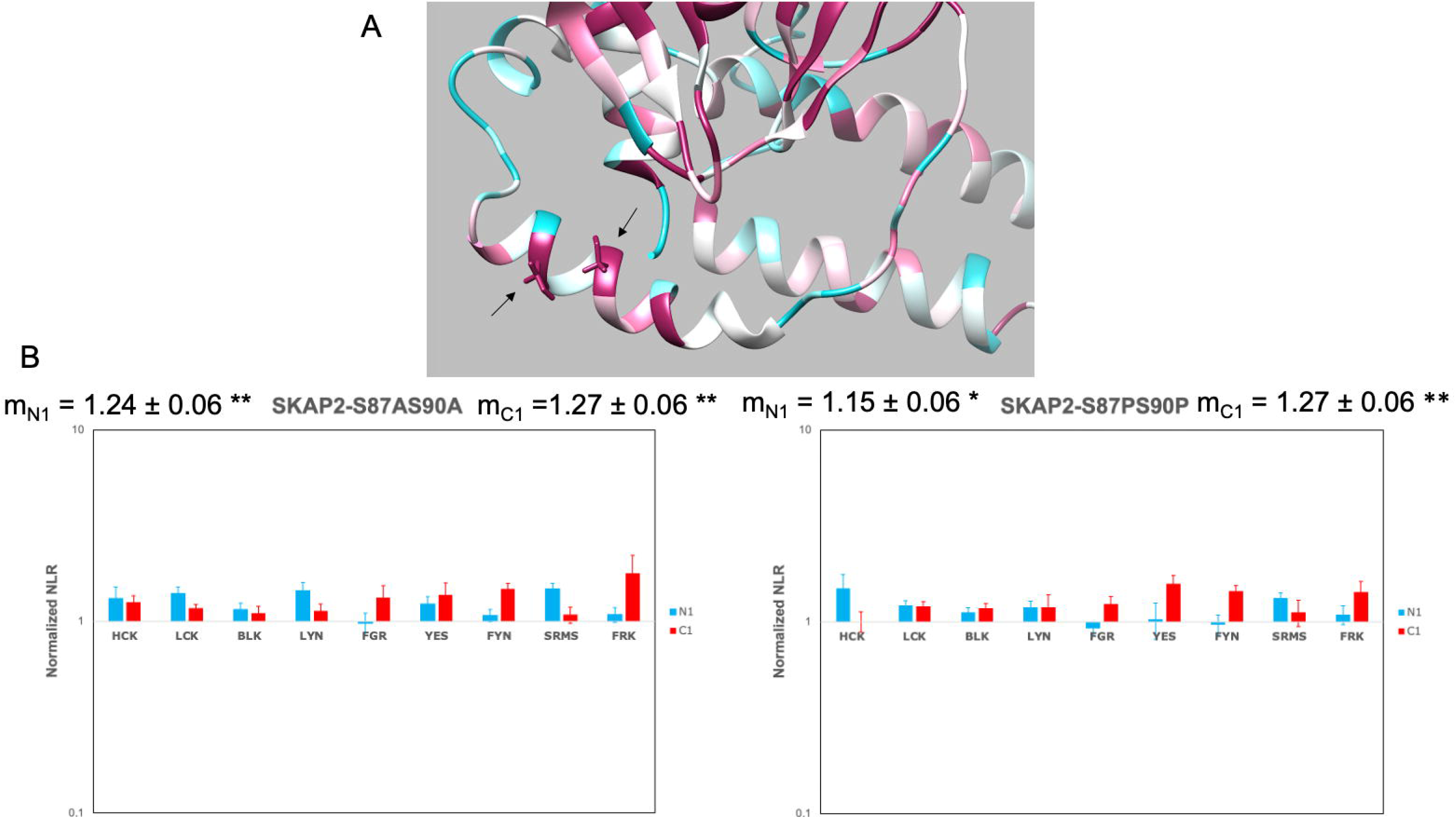
Role of a conserved site of phosphorylation for Casein kinase 1 in Link1 interdomain. (A) The N-terminal part of one chain of SKAP2 from the DIM domain to the PH domain has been modelized by I-Tasser server [66]. Conservation has been evaluated by Consurf website [67, 68] using a scale from highly variable residues (dark blue) to highly conserved one (dark red) passing through white. The phosphorylation site with its α-helix and the two highly conserved serine residues (arrows in the lower left) is in the Link1 interdomain between the DIM domain with its α-helix (lower right) and the PH domain (upper). (B) Graphs show wild type normalized NLR of two SKAP2 mutants, S87AS90A and S87PS90P, with N1-fused (blue) and C1-fused (red) SRC kinases in a logarithmic scale. Wild type normalized NLR is the ratio of the NLR between a SKAP2 mutant and one SRC kinase to the corresponding NLR between wild type SKAP2 and the same SRC kinase fused with hemi-gaussia at the same position. For each graph, the normalized NRL mean ± SEM among SRC kinases is shown as its level of significance (*p* = 0.05 (*), *p* = 0.01 (**), *p* = 0.001 (***)).

### Role of two conserved motifs located in the interdomain Link2

The interdomain Link2 plays a major role for the binding of SRC kinases on both SKAP1 and SKAP2 proteins supporting that similar motifs exist for the two proteins. Search for motifs conserved in both SKAP1 or SKAP2 phylogeny found two motifs consisting of a binding motif for a SH2 domain followed or forward by a glutamine acid-rich region [70, 71]. We decided to test their effect on the binding to SRC kinases by delating one of them or both in the two proteins. Figure 5 shows the result of four experiments for SKAP1 (A) and six experiments for SKAP2 (B) mutants. For SKAP2, we have also tested the mutation of three tyrosine to phenylalanine located at position 75, 237 and 261, which have been predicted to be inside a binding motif for a SH2 domain. This mutant, named 3YF, was mutated at the three predicted binding motifs for a SH2 domain. For each of the two proteins, deletion of one motif slightly decreases the binding to SRC kinases and deletion of both motifs has a greater effect than the deletion of each separatly. Note also that the binding capacity to SRC kinases appears similar or slightly greater for the 3YF SKAP2 mutant than for ΔLink2AB SKAP2 mutant. Supplemental Table 3, which shows the results of the statistical analysis, supports this analysis. Interestingly, the deletion of motif A affects differently the level of the PPI signal than that of motif B depending on SRC kinases. This difference frequently persists in both SKAP1 and SKAP2 proteins. For example, the decrease of binding for HCK, LYN, FYN, and SRMS kinases is greater after deletion of motif A than that of motif B in both proteins and the opposite is true for LCK. But the decrease of binding for BLK and FRK kinases is greater after deletion of motif A than that of motif B only for SKAP2.

**Figure 5:**
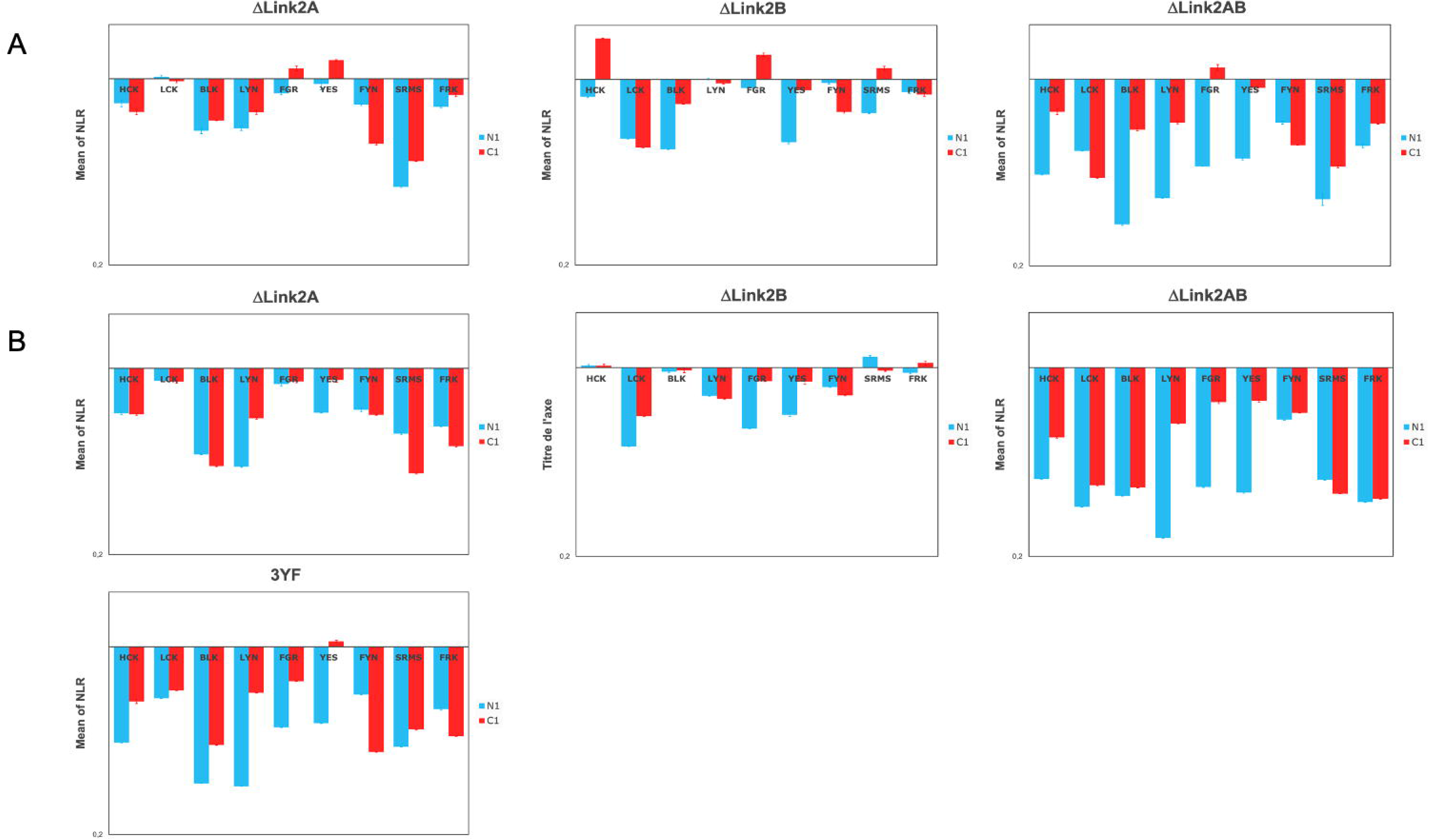
Study of the deletion of the two conserved motif in the interdomain Link2 for SKAP1 and SKAP2 protein. Graphs show wild type normalized NLR of three SKAP1 deletion mutants and four SKAP2 mutants with N1-fused (blue) and C1-fused (red) SRC kinases in a logarithmic scale. Wild type normalized NLR is the ratio of the NLR between a mutant and one SRC kinase to the corresponding NLR between wild type protein and the same SRC kinase fused with hemi-gaussia at the same position. ΔLink2A: deletion of motif A; ΔLink2B: deletion of motif B; ΔLink2AB: deletion of both motifs; 3YF: mutation to phenylalanine of the three tyrosine residues located at position 75, 237, and 261 of SKAP2.

### Dissociating the binding capacity of the SH2 domain from activation for HCK and LCK kinase

These two conserved motifs contain a binding site for SH2 which role for binding to SRC kinases was evaluated. It is not a trivial task because the SH2 domain of SRC kinases also plays a major role for their activation (Fig. 1C). To dissociate the activation to the binding capacity of the SH2 domain of the kinase, we decided to study two mutants, either Y501F and R150A for HCK kinase or Y505F and R154A for LCK one. The R150A HCK/ R154A LCK mutant has lost the binding capacity of the SH2 domain and only activation of HCK/LCK kinase occurs (Figure 1C). The Y501F HCK/ Y505F LCK mutant has lost the binding site of its SH2 domain and activation of the HCK/LCK kinase occurs associated to the capacity of its SH2 domain to interact with other proteins. To dissociate a direct effect of SKAP1 and SKAP2 to an indirect effect of other proteins, which binding modified the interaction with SKAP1 and SKAP2, we also study their ΔLink2AB mutant. The indirect role is detected by a difference of ΔLink2AB binding between the mutant bearing the binding motif to SH2 domain and that of the SH2 binding site and the direct role by a decrease of ΔLink2AB binding compare to wild type adaptor with mutant bearing the binding motif to SH2 domain that disappears with mutant of the SH2 binding site. Figure 6 shows the result of one experiment for SKAP2 (upper) or SKAP1 (lower) challanged against HCK (A) or LCK(B) mutants. First, we will analyze HCK data (Fig. 6A). For SKAP2 (upper), the activation of the HCK kinase shown by comparing results of wildtype HCK to R150A mutant increases the interaction between SKAP2 and HCK independently of the localization of the hemi-gaussia fusion. The role of the SH2 domain obtained by comparing results of Y501F and R150A HCK mutants shows that its effect on the binding to SKAP2 is moderate (N1-fused data) or null (C1-fused data). Study of ΔLink2AB mutant confirms that results and suggests that the moderate effect shown in N1-fused Y501F data is due to an indirect effect. The study of 3YF SKAP2 mutant suggests also that the glutamic acid-rich region plays a role since differences of signal between ΔLink2AB mutant and 3YF one was present for wildtype HCK and N1-fused Y501F HCK mutant. However, we cannot exclude a role of Y75F mutation. Similar analysis for SKAP1 (Fig. 6A lower) shows different results. If activation of HCK has a similar effect than for SKAP2, the role of the SH2 domain was completely different. The difference of PPI signal between wildtype SKAP1 and ΔLink2AB mutant suggests that both a direct and an indirect effect of the SH2 domain occur. Note that the results for ΔPHLink2 mutant is completely different supporting that the PH domain and/or the interdomain Link2 play other role in this interaction.

**Figure 6:**
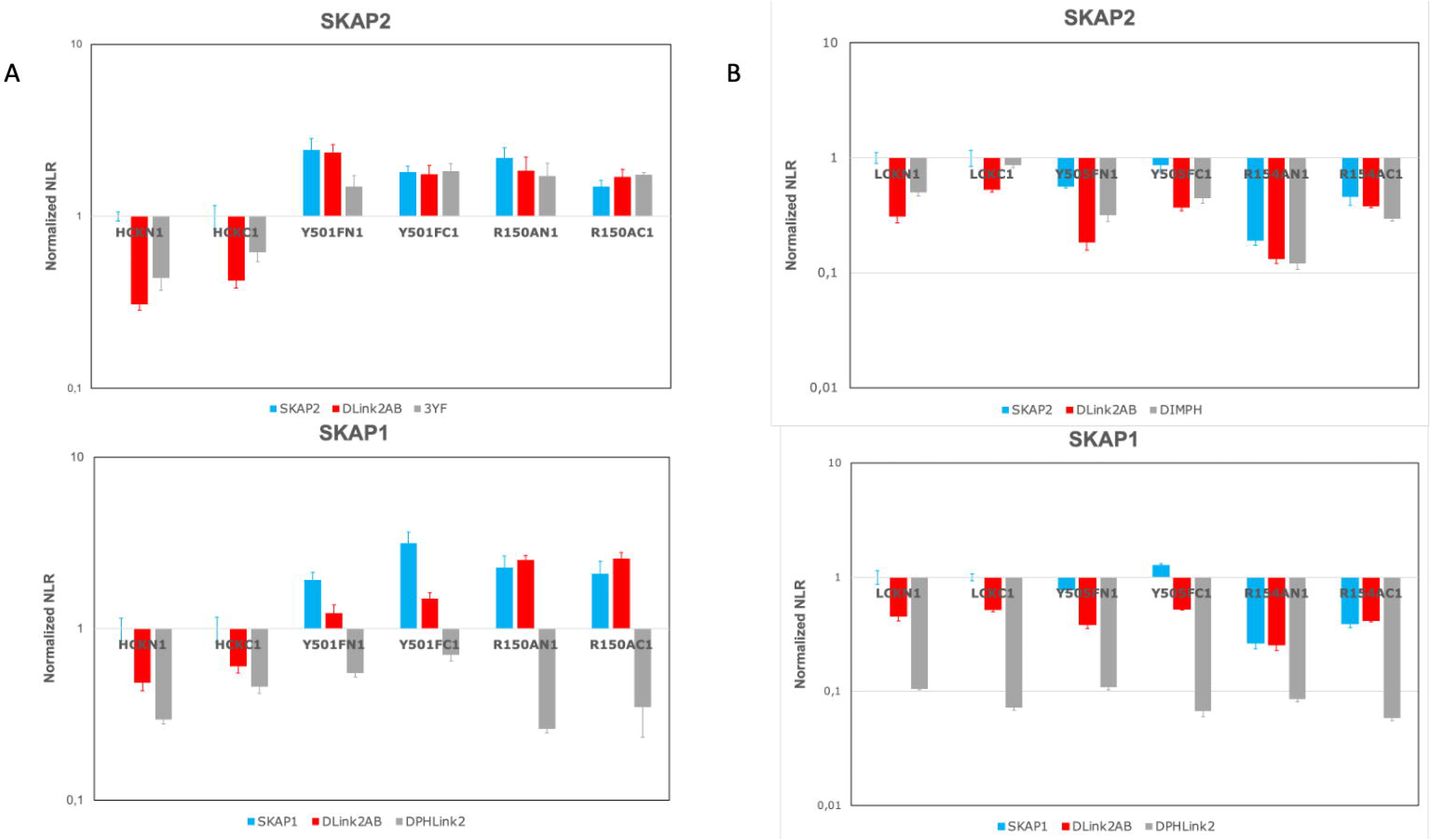
Dissociating the role of activation and inactivation of its SH2 domain in the interaction with SKAP1 and SKAP2. Graphs show wild type normalized NLR of two SKAP2 mutants (upper) and two SKAP1 deletion mutants (lower) with either N1-fused and C1-fused wild type HCK and HCK mutants (A) or N1-fused and C1-fused wild type LCK and LCK mutants (B) in a logarithmic scale. Wild type normalized NLR is the ratio of the NLR between a couple of SKAP2 and kinase mutants and the corresponding NLR between wild type SKAP2 and wild type kinase fused with hemi-gaussia at the same position. Similar normalization was performed for SKAP1 data. ΔLink2AB: deletion of motif A and B in interdomain Link2; 3YF: mutation to phenylalanine of the three tyrosine residues located at position 75, 237, and 261 of SKAP2; ΔPHLink2: deletion of the PH domain and interdomain Link2: DIMPH: N-terminal part of SKAP2 containing the DIM and the PH domains; Y501F HCK and Y505F LCK mutations inactivate the intramolecular binding to the SH2 domain in contrast to R150A HCK and R154A LCK mutations, which inactivate the binding capacity of SH2 domain to its phosphorylated motif.

Analyzing LCK data (Fig 6B) shows a completely different situation than for HCK data. For both SKAP2 (upper) and SKAP1 (lower), LCK activation decrease the PPI signal. This difference between the two SRC kinases are probably due to a difference of binding for the second module since DIMPH SKAP2 mutant has a similar pattern than wild-type SKAP2. A direct role of the SH2 domain is clearly seen for SKAP1 and also SKAP2 even if an indirect role is possible for both.

In conclusion, this analysis supports a strong increase of HCK binding due to its activation for both SKAP1 and SKAP2 proteins in contrast to LCK where the opposite is true. The SH2 domain of HCK plays a minor role for the interaction with SKAP2 in contrast to SKAP1. For LCK, its SH2 domain seems to play a similar role for both adaptors. Interestingly, study of DIMPH SKAP2 mutant suggests that the decrease of binding of LCK after its activation is due to the DIM domain. As previously mentioned, the study of 3YF SKAP2 mutant suggests that the two conserved motifs contain more than a unique site for SH2 domain.

### Study of the three FYN isoforms

The opposite effect of activation on the binding capacities between HCK and LCK kinases suggests studying the three FYN isoforms. Figure 7 shows the result of this experiment using as reference N1-fused and C1-fused wild type FYN isoform. The pattern of wild type isoforms and their ΔLink2AB mutants is similar for SKAP1 and SKAP2 and will be described together. Wild type FYNT has a lower PPI signal in contrast to wild type FYNB, which has an upper signal. The ΔLink2AB mutants have a similar PPI signal than their wild type isoforms for C1 data in contrast to N1 data where this signal is lower. DIMPH SKAP2 mutant shows a similar pattern supporting than the PPI signal of the wild type isoform depends mainly of the second module and the DIM domain. For N1-fused data, an effect of the two conserved motifs is also detected. As previously reported, the ΔPHLink2 SKAP1 mutant has a different pattern than the ΔLink2AB SKAP1 mutant supporting than other factors than the two conserved motifs influence the PPI signal. The different binding capacity of these isoforms are due to exon 7 coding for the C-terminal part of the SH2 domain (fourteen last aa), the hinge, and the N-terminal part of the tyrosine-kinase catalytic domain (seventeen first aa).

**Figure 7:**
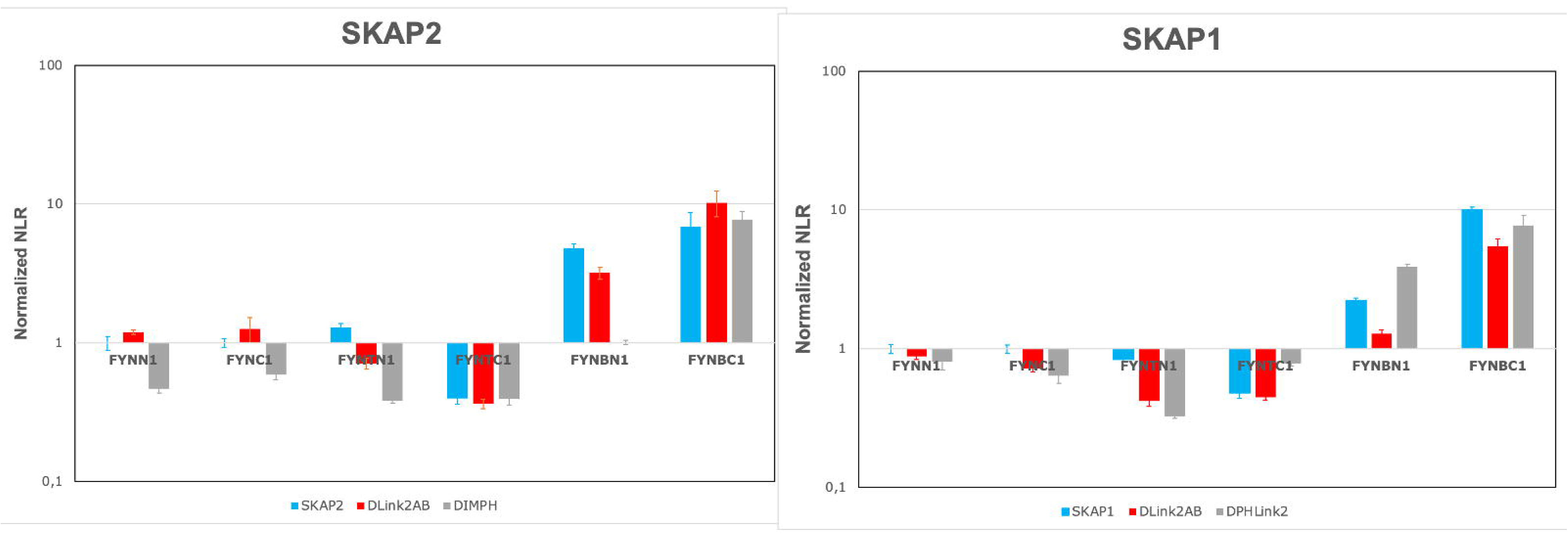
The level of the PPI signal with SKAP1 and SKAP2 is lower for FYNT isoform than for FYNB isoform. Graphs show FYN normalized NLR of SKAP2 (left) and SKAP1 (right) for the three wild type FYN isoforms and their mutants. N1 data are normalized according to N1-fused wild type FYN and C1 data according to C1-fused wild type FYN. See Figure 6 caption for details.

## Discussion

The characterization of the SKAP1 modular architecture, which recognizes SRC kinases allowed us to determine that (1) two modules control these interactions; (2) each module has a different effect depending on SKAP1 and SKAP2 [27]. If a general pattern might be obtained for all SRC kinases, each one has also its own characteristics. These results illustrate the interest of luciferase complementation assay to dissect how proteins of the same family interact with other proteins. Understanding how evolution shapes such interactions has important consequences.

By comparing the present results to previous ones on SKAP2 [27], we showed that both SKAP1 and SKAP2 modular architectures are globally similar, but exhibit differences mainly between C1-fused SRC kinases data and N1-fused ones (supplemental Figure S4 and [27]). The DIM domain of SKAP2 alone binds efficiently some C1-fused SRC kinases in contrast to SKAP1. In the present work, we confirmed these results showing that both SKAP1 modules, the DIM domain and the Link2 interdomain, are necessary to bind SRC kinases independently of the fusion localization. We also showed that the SH3 domain of SKAP1 does not have a global effect on these interactions in contrast to that of SKAP2 [27]. In both proteins, this SH3 domain has an indirect effect due to its binding capacity to interactors, which probably affects the localization of these adaptors and their interaction with SRC kinases. In the present paper, we showed also that each conserved motif located in the Link2 interdomain recognizes differently SRC kinases with a conserved pattern between SKAP1 and SKAP2 (Figure 5). A principal component analysis [65] confirmed that two different patterns due to either the SH3 domain or the two motifs in the Link2 interdomain, are detected on this C1-fused kinases data of SKAP2 (Supplemental data). A similar result was obtained on the C1-fused kinases data of SKAP1, supporting that the pattern due to the SH3 domain is not lost in SKAP1. It has not been detected only because its global effect is small or null. Note that this result explains that the pattern of ΔSH3 SKAP1 mutant is so different from that of the W333K SKAP1 mutant even if there global effect is similar (Figure 3). However, each deletion of one of the two conserved motifs located in Link2 interdomain proteins induces a similar pattern between N1-fused and C1-fused SRC kinases in both SKAP1 and SKAP2 (Figure 5 and principal component analysis). These results support that for both proteins each module recognizes differently each kinase independently to the other module and that these patterns are globally conserved since their duplication.

To find out more, search for conserved motifs in the interdomain Link2 common to both proteins allows to define two motifs composed of an SH2 binding site followed or forwards by a glutamic-acid rich region [70]. These two motifs play non-redundant and complementary role for binding to SRC kinases in both proteins (Figure 5). Interestingly, each of these motifs recognizes similarly some SRC kinases (HCK, FYN, LCK, LYN, and SRMS) for both adaptors. There are some exceptions as BLK and FRK, which are specifically recognized by motif A of SKAP2. These results support that such evolutionary conservation of two motifs for both proteins is partially due to their non-redundant role. A second possible mechanism may be a cooperative effect between the two motifs. Interestingly, ΔLink2AB deletion induces a greater decrease of the binding to SRC kinases than the addition of ΔLink2A and ΔLink2B effect at least for N1-fused SRC kinases with both SKAP1 and SKAP2 (Supplemental Table S3). Motifs recognized by SH2 domain have been subject to a lot of work such as those for SRC kinases [72, 73]. The two motifs in Link2 interdomain are characteristic of a binding motif to their SH2 domain even if motif B is the more conformed. In our knowledge, a glutamic acid rich region has not been associated to such interactions. Note also, that other putative signals such as a Y based sorting signal or an Atg8 protein family binding site overlapped these two regions in which tyrosine residues should be unphosphorylated [70]. We cannot exclude that these motifs play an indirect role. Lastly, we showed that the interdomain Link1 affects these interactions only in the case of SKAP2 (Figure 3 and Table 3). Search for a conserved motif specific to SKAP2 was able to detect a conserved phosphorylation site for Casein kinase 1, which deletion increases the interaction with SRC kinases by a direct or indirect effect (Figure 4).

The respective effect of the binding capacity of the SH2 domain and kinase activation was studied for HCK and LCK kinases interacting either with SKAP1 or SKAP2 (Figure 6). In most cases, a direct effect of the SH2 domain was detected except for SKAP2 interacting with the HCK kinase. In constrast, an opposite effect of kinase activation was observed for HCK and LCK kinases. Interestingly, kinase activation increased the binding of HCK kinase, which is mainly expressed in monocytes/macrophages lineage and neutrophils as SKAP2, in which the second module and the DIM domain play a higher role than in SKAP1. In contrast, kinase activation decreased the binding of LCK kinase, which is mainly expressed in T lymphocytes as SKAP1. This suggests that 1) the effect of kinase activation mainly depends on the second module and the DIM domain 2) a convergent evolution between SRC kinases and SKAP1 or SKAP2 might occur. The first part of the hypothesis is supported by previous data showing that the activation of some SRC kinases increases their binding to SKAP2 and to DIM domains even if the subcellular localization also plays a role [27]. A similar pattern between DIMPH SKAP2 mutant and wild type SKAP2 interacting with wild type LCK and LCK mutants (Figure 6) also supports this hypothesis. The hypothesis of convergent evolution suggests that the binding capacity of the DIM domain alone as in SKAP2 is associated with an increase of binding by kinase activation. This property is absent for kinases interacting with SKAP1 as suggested by the study of LCK. Study of the three FYN isoforms supports this hypothesis (Figure 7): FYNT isoform expressed in T lymphocytes as SKAP1 has its binding to SKAP1 and SKAP2 decreasing compare to FYN isoform in contrast to FYNB isoform expressed in the brain as SKAP2. Study of ΔLink2AB mustants and of DIMPH SKAP2 mutant agrees with that hypothesis. Direct interactions between SKAP2 and HCK in monocytes/macrophages lineage [10] or FYN in the brain [9, 74] have been reported in the litterature as well as between SKAP1 and FYN [4, 14, 16] in T lymphocytes. However, this hypothesis has to be confirmed by further works.

In conclusion, this work dissected the SKAP1 modular organization recognizing SRC kinases defining two modules common also to SKAP2 modular organization but with different properties. Comparison with that of SKAP2 allowed us to define the role of different conserved motifs in these interactions, which had interesting consequences on how evolution shapes these protein-protein interaction. Lastly, study of HCK and LCK kinases as well as FYN isoforms suggests an interesting hypothesis of coevolution between the properties of these kinases and those of SKAP1 and SKAP2 adaptors.

## Acknowledgements

We thank Patricia Cassonnet, and Mélanie Dos Santos for technical supports, Chen Kuang-Yu, Tim Krischuns and Guillaume Dugied for regularly providing HEK 293T cell culture, and the Dana-Farber Cancer Institute (Boston, MA, USA) for providing us with ORFs.

## Availability of data and materials

The datasets used and/or analyzed during the current study as well as mutants generated in the present study are available from the corresponding author on reasonable request.

## Ethics approval and consent to participate

This research has received an OGM agreement from the Ministère de l’Education Nationale (Dossier DUO n°9216).

## Competing interests

The authors declare no competing interests.

## Funding

Funding was provided by Institut Pasteur (Paris), Centre National de la Recherche Scientifique, Université Paris Diderot, Sorbonne Paris Cité.

## Author contributions

**Laurine Levillayer:** Investigation **Camille Brighelli:** Investigation **Caroline Demeret:** Writing - Review & Editing **Anavaj Sakuntabhai:** Funding acquisition **Jean-François Bureau:** Conceptualization, Formal analysis, Investigation, Writing - Original Draft

## Supporting Informations

**Supplemental Tables** An excel file containing S1 Table Sequence of new primers used in this study; S2 Table Statistical analysis of deletion of motif A, and B alone or in combination in SKAP1 (upper part) and SKAP2 (lower part); S3 Table Search for an cooperative effect between motif A and B.

**Supplemental data** A docx file containing the four supplemental figures and the principal component analysis.

## Notes

### Competing Interest Statement

The authors have declared no competing interest.

