## Supplemental_data for "Role of two modules controlling the interaction between SKAP1 and SRC kinases Comparison with SKAP2 architecture and consequences for evolution"

**Supplemental figures**

**Figure S1: Effect of SKAP1 mutants on the interaction with SRC kinases.** Differential interaction scatterplot of SKAP1 mutant (y-axis) versus SKAP1 wild-type (x-axis) using NLR are displayed in (A-F). Interactions not affected by the mutation are aligned on the diagonal. PPI disruptive mutations are in the lower right-hand quadrant in contrast to mutations stabilizing interaction that are located on the upper left-hand one. Interactions for N1-fused SRC kinase are in blue circle and those for C1-fused SRC kinase are in red square. Error bar: Standard Error to the Mean (SEM). A robust linear regression, which takes account of outliers, was performed on data using mmregress Stata module. Linear regression equations are (A) log(ΔPHLink2) = 1.209 * log(SKAP1) – 0.750 for C1-fused SRC kinases; (B) log(DomDIM) = 0.970 * log(SKAP1) - 0.985 for N1-fused SRC kinases and log(DomDIM) = 0.600 * log(SKAP1) + 0.112 for C1-fused SRC kinases; (C) log(ΔSH3) = 0.912 * log(SKAP1) + 0.0562 for N1-fused SRC kinases; (D) log(ΔLink1PH) = 1.070 * log(SKAP1) + 0.043 for N1-fused SRC kinases and log(ΔLink1PH) = 1.115 * log(SKAP1) - 0.122 for C1-fused SRC kinases; (E) log(D120K) = 1.375 * log(SKAP1) – 0.691 for N1-fused SRC kinases and log(D120K) = 1.030 * log(SKAP1) – 0.042 for C1-fused SRC kinases; (F) log(W333K) = 0.980 * log(SKAP1) + 0.120 for N1-fused SRC kinases and log(W333K) = 0.983 * log(SKAP1) + 0.123.

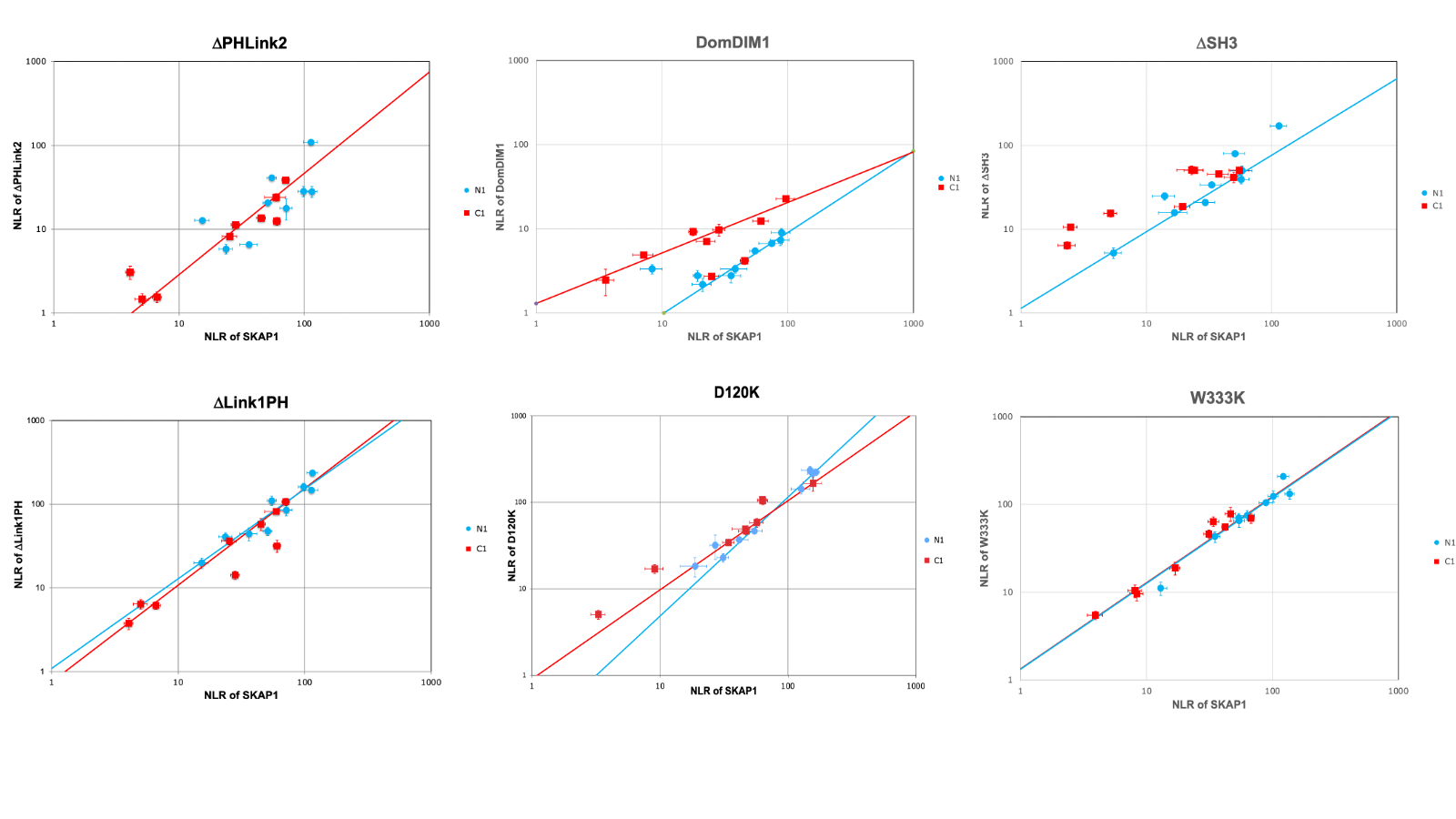

**Figure S2: Comparison of mutations between SKAP1and SKAP2 on the interaction with SRC kinases.** Differential interaction scatterplot of SKAP1 mutant (y-axis) versus the same SKAP2 mutant (x-axis) using NLR are displayed in (A-J). Mutants in which SKAP1 and SKAP2 play similar role are aligned on the diagonal. Mutants in which SKAP1interacts with higher efficiency to SRC kinases than SKAP2 are in the lower right-hand quadrant in contrast to mutants in which SKAP1interacts with lower efficiency to SRC kinases than SKAP2 that are located on the upper left-hand one. Interactions for N1-fused SRC kinase are in blue circle and those for C1-fused SRC kinase are in red square. Error bar: Standard Error to the Mean (SEM). Stars are the barycenter of each cloud. A classical linear regression was performed on data using regress Stata module. Linear regression equations are (A) log(SKAP1) = 0.755 log(SKAP2) + 0.0354 for N1-fused SRC kinases and log(SKAP1) = 0.964 log(SKAP2) -0.404 for C1-fused SRC kinases; (B) log(D120K) = 0.684 * log(D129K) + 0.456 for N1-fused SRC kinases and log(D120K) = 0.884 log(D129K) -0.225 for C1-fused SRC kinases; (C) log(DIM-) = 0.633 log(DIM-) + 0.622 for N1-fused SRC kinases and log(DIM-) = 0.965 log(DIM-) – 0.085 for C1-fused SRC kinases; (D) log(ΔLink1PH) = 0.776 * log(ΔLink1PH) + 0.686 for N1-fused SRC kinases and log(ΔLink1PH) = 1.180 * log(ΔLink1PH) -0.296 for C1-fused SRC kinases; (E) log(ΔPH) = 0.977 * log(ΔPH) – 0.146 for N1-fused SRC kinases and log(ΔPH) = 0.906 * log(ΔPH) – 0.447 for C1-fused SRC kinases; (F) log(ΔPHLink2) = 1.009 * log(ΔPHLink2) – 0.144 for N1-fused SRC kinases and log(ΔPHLink2) = 0.869 * log(ΔPHLink2) – 0.329 for C1-fused SRC kinases; (G) log(ΔSH3) = 1.026 * log(ΔSH3) + 0.081 for C1-fused SRC kinases; (I) log(Link2SH3) = 1.110 * log(Link2SH3) – 0.235 for C1-fused SRC kinases; (J) log(W333K) = 0.841 * log(W336K) + 0.306 for N1-fused SRC kinases and log(W333K) = 1.021 * log(W336K) – 0.355 for C1-fused SRC kinases.

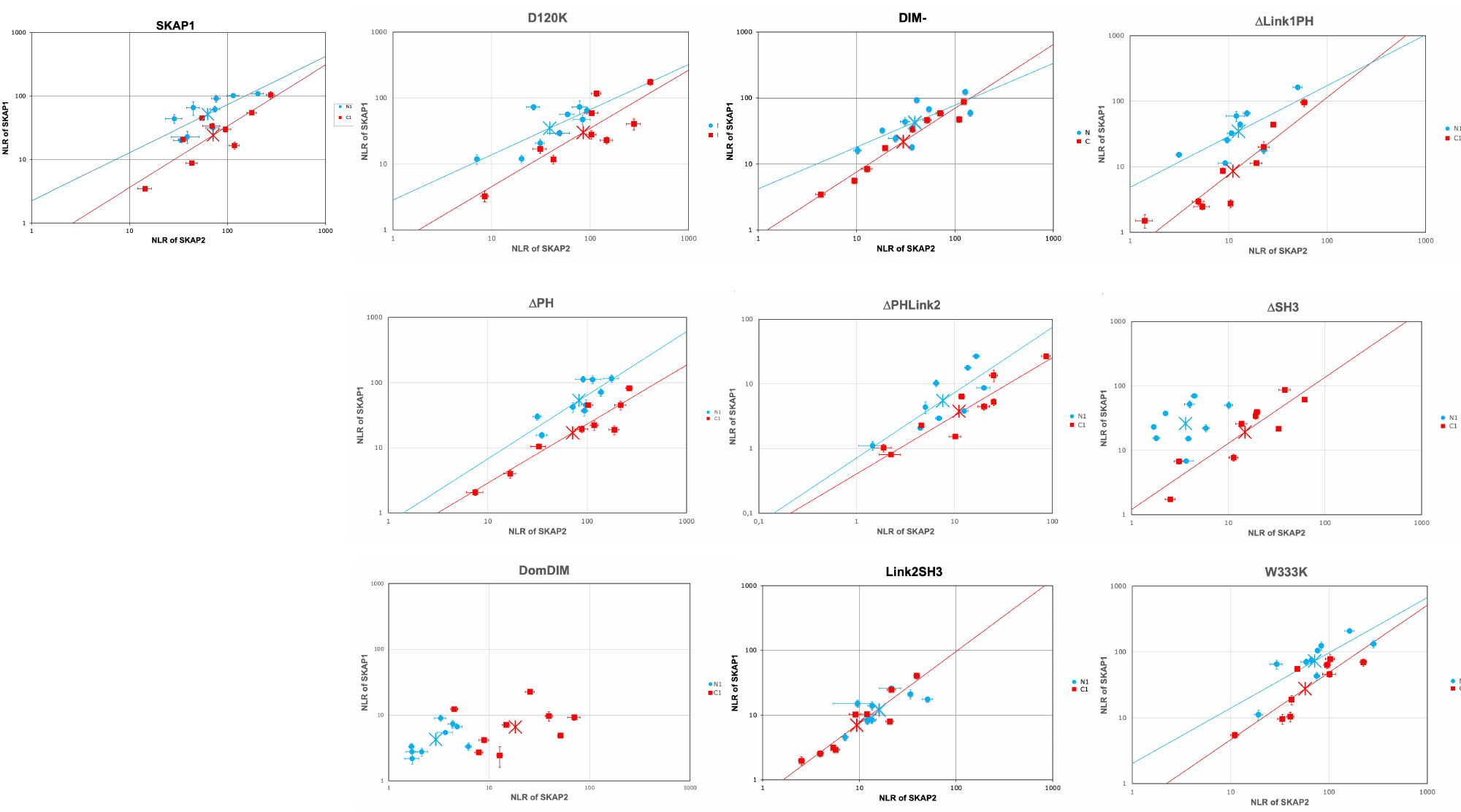

**Figure S3: Scatterplot of the barycenters between SKAP1 and SKAP2 data.** Barycenters from each of the ten wildtype or mutants of figure S2 were plotted. Mutants in which SKAP1 and SKAP2 play similar role are aligned with the wild type adaptors. Mutants in which SKAP1interacts with higher efficiency to SRC kinases than SKAP2 are in the lower right-hand quadrant in contrast to mutants in which SKAP1interacts with lower efficiency to SRC kinases than SKAP2 that are located on the upper left-hand one. Interactions for N1-fused SRC kinase are in blue circle and those for C1-fused SRC kinase are in red square. Barycenter of two experiments comparing wildtype SKAP1 to wildtype SKAP2 are shown in darker colors. The name of mutants in which SKAP1 and SKAP2 play different roles are indicated.

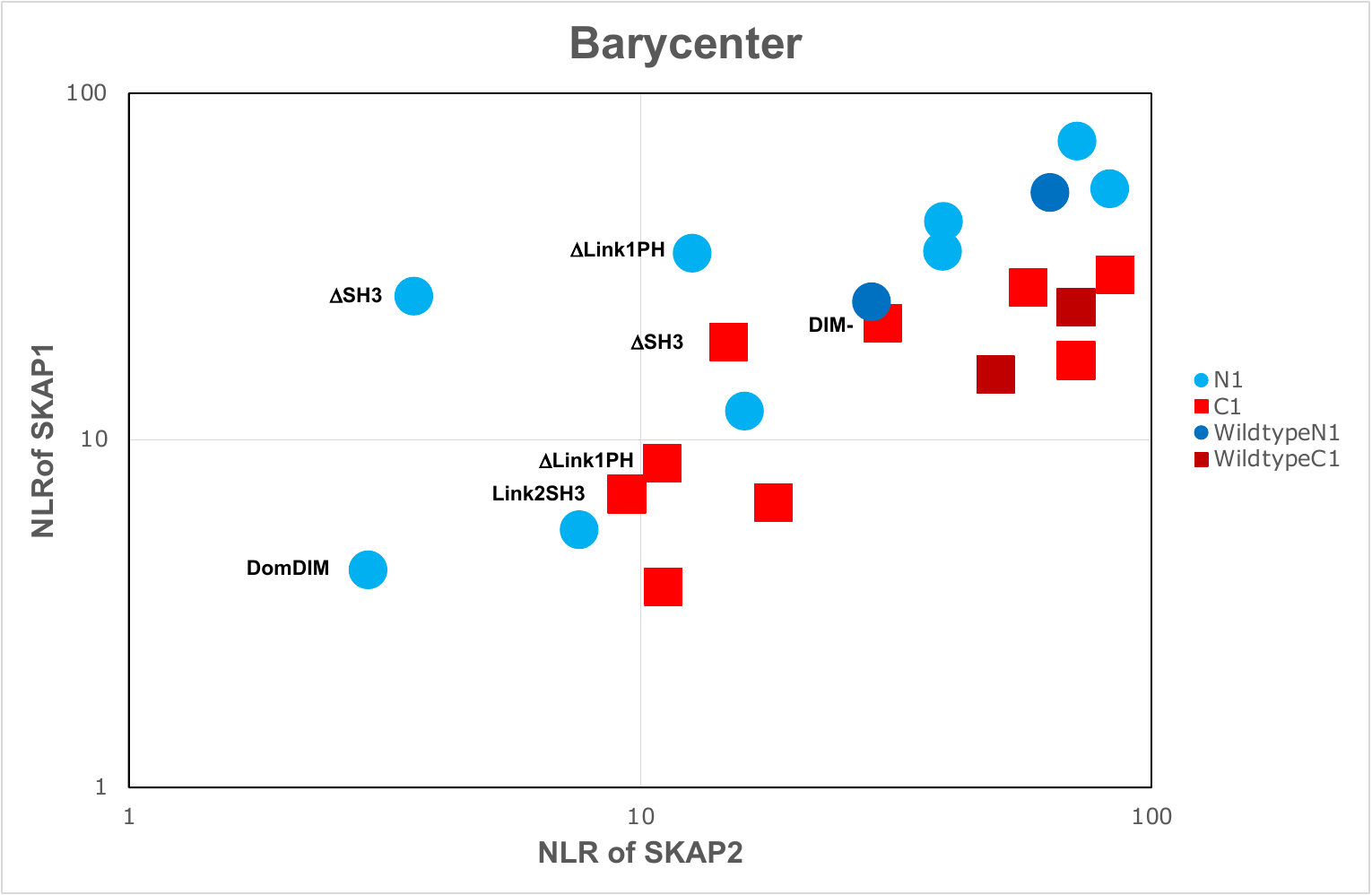

**Figure S4: means of NLR for N1 or C1-fused wild-type SRC kinases.** Data of the 26 experiments with SKAP1 mutants were used to calculate for each of the nine wild type SRC kinases either N1-fused (blue) or C1-fused (red) the mean of the NLR ± SEM.

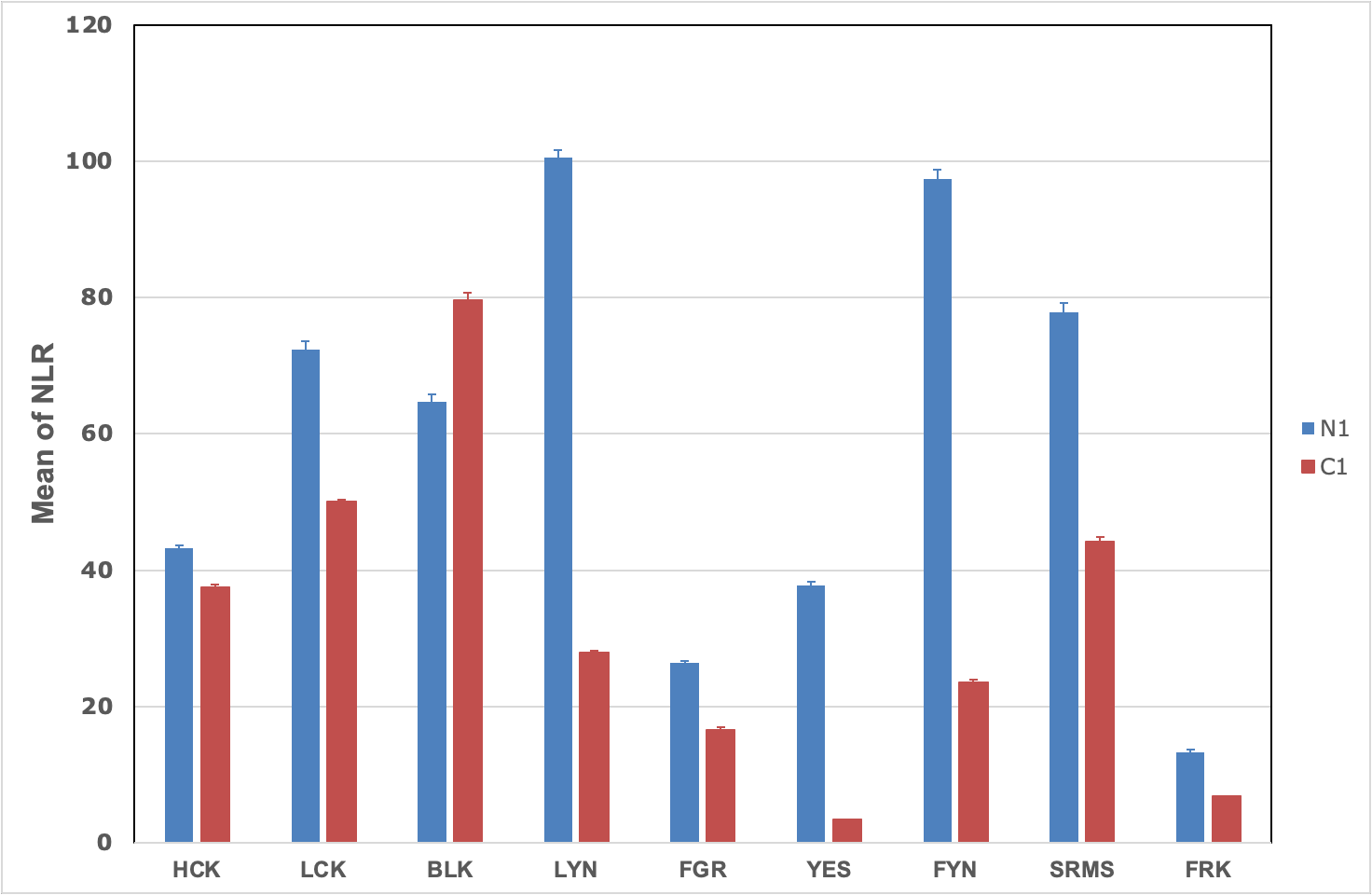

**Principal component analysis**

A principal component analysis (PCA) was performed on SKAP1 and SKAP2 mutant data to determine which factors explained the variability amongst C1-fused SRC kinases and to compare them between the two paralogs.

Methodology

A PCA on SKAP2 mutant data from our previous paper (1) in which was added those concerning deletion of motifs A and B was performed after a log-transformation using the R packages FactomineR (2) and Factoshiny and a second one on SKAP1 mutant data from the present paper. The mean value from at least three experiments of the log-transformed normalized NLR of each mutant and each C1-fused SRC kinase was centered and reduced depending on the nine SRC kinase data. A qualitative variable defining the type of mutation was added as a supplemental non explicative variable with four modalities: deletion mutation (Del), nonsynonymous mutation alone (Mut) or multiple (Mut2), deletion mutation associated to the study of motif A and B (Exp). We used the decomposition of the total inertia to determine the number of dimensions to analyze. We analyzed the graph of mutants and those of the C1-fused SRC kinases. After the PCA, a hierarchical clustering (HC) was performed followed by a consolidation using K mean method in which the number of groups was determined from the PCA. The hierarchical clustering tree and the factor map are analyzed.

Results and discussion

Some part of these results, which have been obtained from the files automatically generated during the PCA and HC analyses. appear in italic.

Analysis of SKAP2 data

No outlier was detected. which could bias the PCA.

Figure 1: Decomposition of the total inertia from SKAP2 data

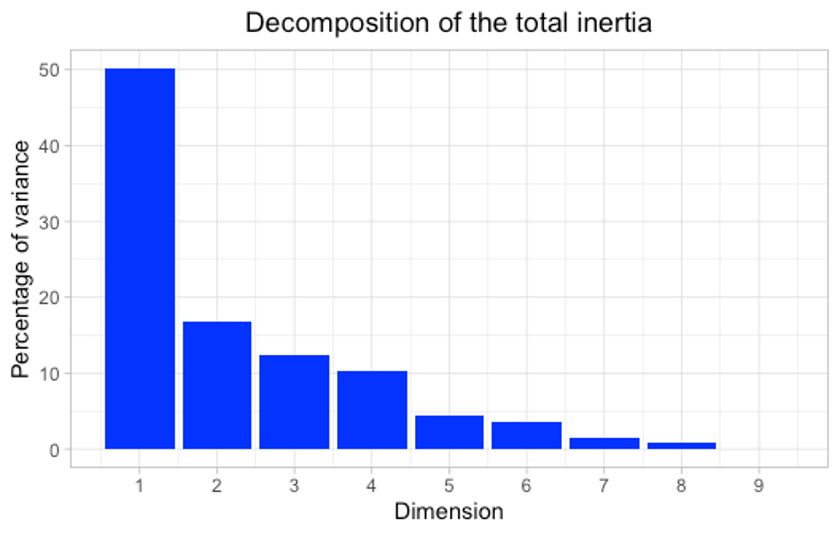

*The first two dimensions express* ***67%*** *of the total dataset inertia. This percentage is relatively high and thus the first plane well represents the data variability. This value is greater than the reference value that equals* ***47.2%*** *(3). the variability explained by this plane is thus significant (the reference value is the 0.95-quantile of the inertia percentages distribution obtained by simulating 11419 data tables of equivalent size based on a normal distribution). The first factor is major: it expresses itself 50.18% of the data variability.* *Note that in such a case. the variability related to the other components might be meaningless. despite of a high percentage.*

Figure 2: Individual factor map of SKAP2 (A) and SKAP1 (B) data. Mutants, in which their color depends on their contribution to the plane construction (point) and barycenter of each type of mutations (square) are presented.

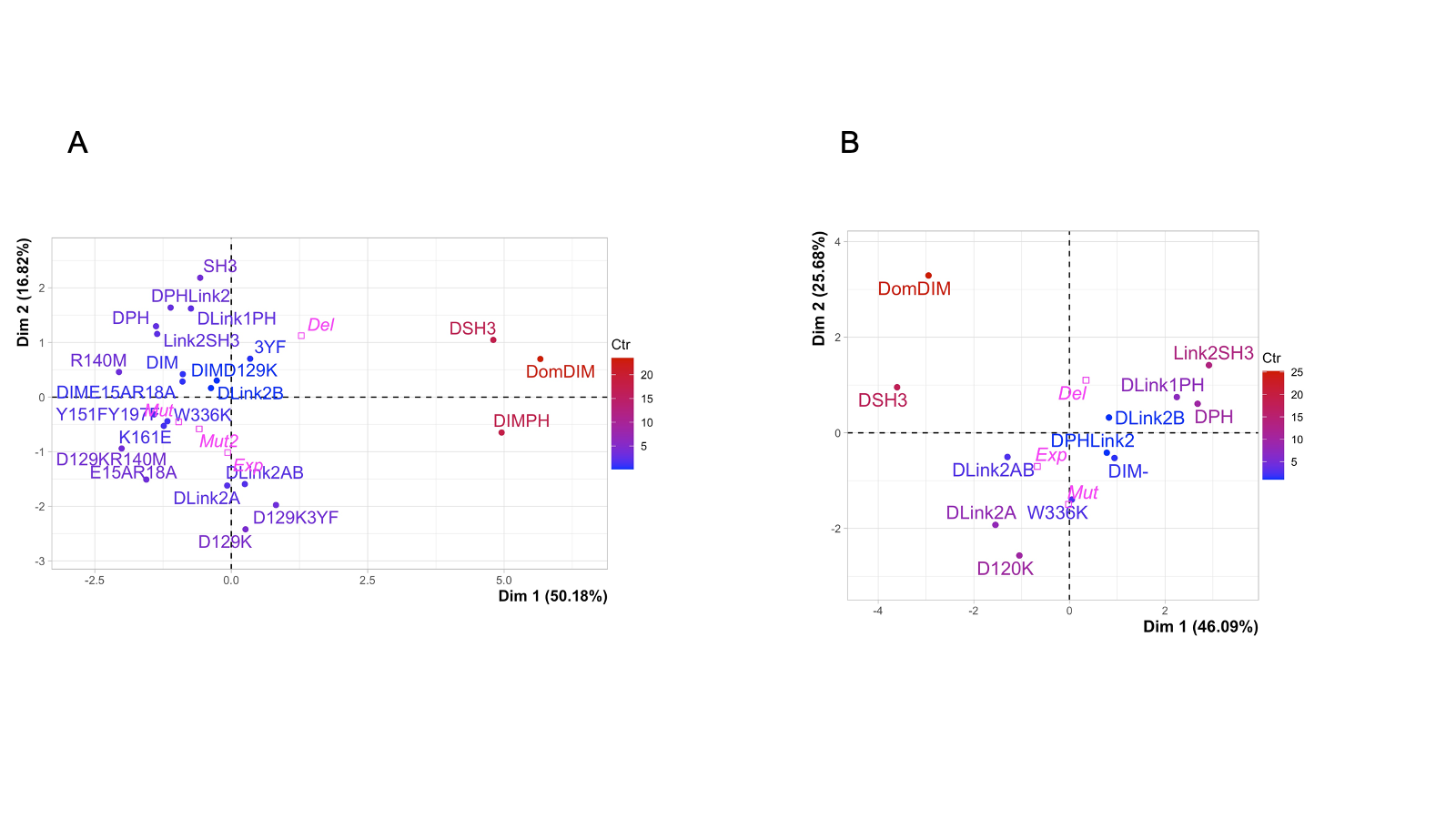

In Figure 2A, the first axis of the plan is well explained by the effect of the SH3 domain. The three mutants, DomDIM, DIMPH, and ΔSH3, in which the DIM domain is present and the SH3 domain is deleted, have the highest value in contrast to all other mutants having low similar values. The second axis explaining only 16.8 percent of the variability amongst SRC kinases, is partially explained by mutants deleted of motif A, B or both. ΔLink2B mutant has middle value in contrast to ΔLink2A mutant, which has one of the lowest values as ΔLink2AB one. This analysis confirmed that the two factors we have previously described during the analysis of SKAP2 data explained an important part of the variability of normalized NLR amongst SRC kinases.

Figure 3 Variable factor map of SKAP2 (A) and SKAP1 (B) data. The position inside the circle of each of the nine SRC kinases is shown. The closest to the circle the position of a kinase is, the more the kinase will be explicative.

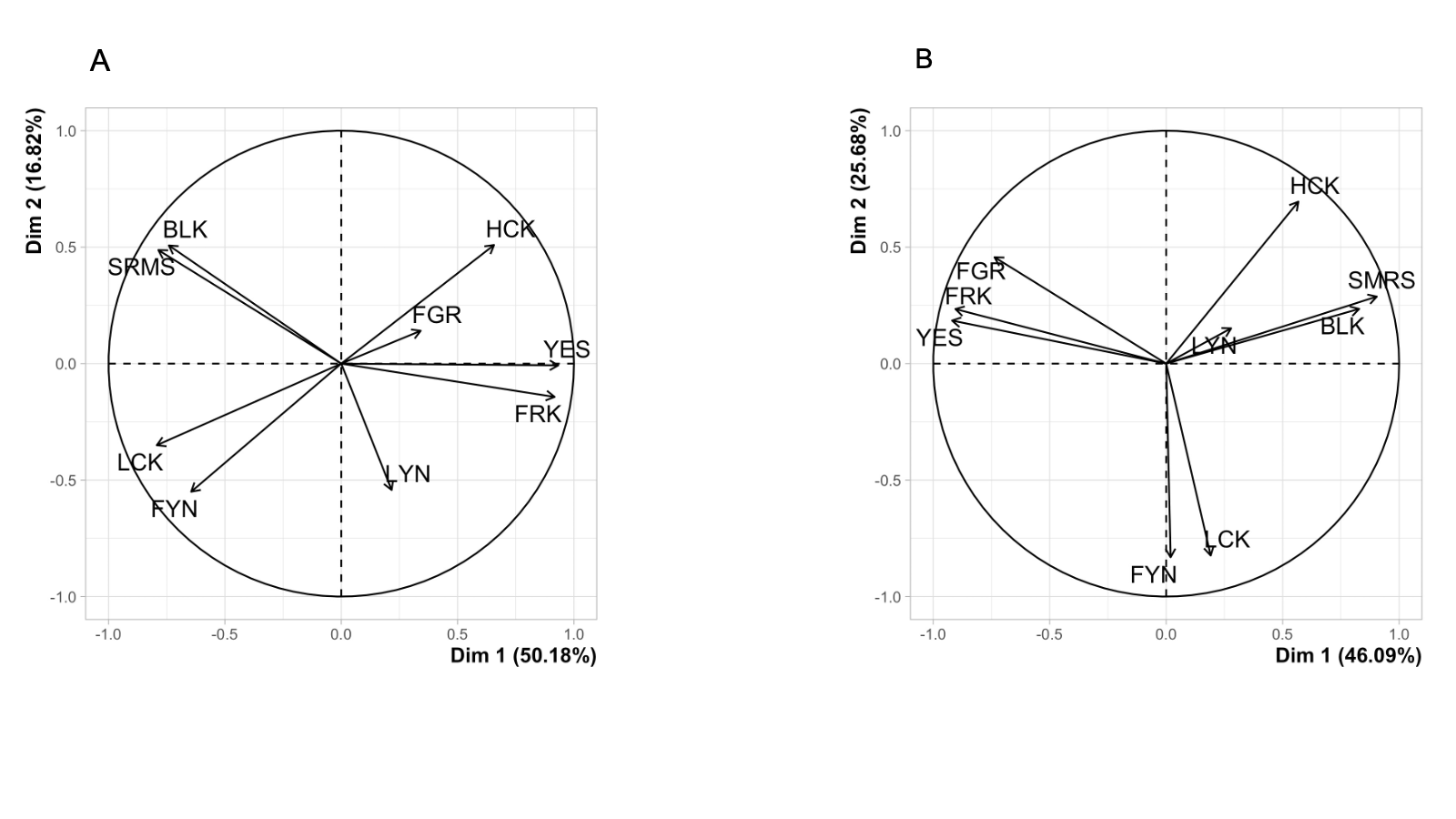

In figure 3A, two kinases, YES and FRK, are positively associated to high values of the first axis in agreement with the highest values of normalized NLR for these two kinases and the three mutants, DomDIM, DIMPH, and ΔSH3 (1).

In conclusion, the two SRC kinase patterns described previously explain a great part of the variability amongst C1-fused SRC kinases in the present PCA of SKAP2 data. This result suggests performing a similar analysis for SKAP1 data to test if similar patterns are detected. Our results on deletion of motif A and/or B suggest that at least this pattern will be found.

Table 1 shows a summary of the data.

Analysis of SKAP1 data

Figure 4 Decomposition of the total inertia from SKAP1 data

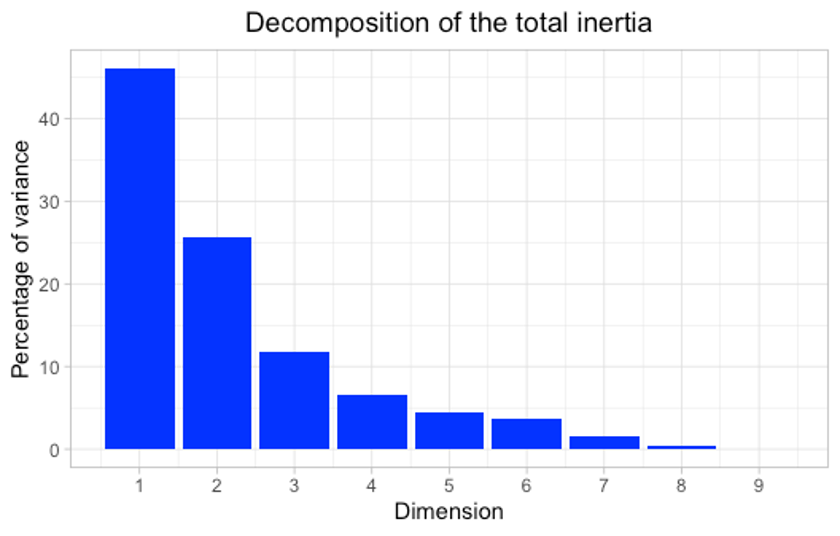

No outlier was detected, which could bias the PCA.

In figure 4, *the first two dimensions express* ***71.77%*** *of the total dataset inertia. This percentage is high and thus the first plane represents an important part of the data variability. This value is greater than the reference value that equals* ***58.47%*** *(3), the variability explained by this plane is thus significant (the reference value is the 0.95-quantile of the inertia percentages distribution obtained by simulating 9397 data tables of equivalent size based on a normal distribution).*

Figure 2B is very similar to Figure 2A even if the effect of the two first axis is more difficult to explain. One of the most important differences is the lowest inertia explains by the first axis. However, the effect of their two diagonals can be explain similarly. A first diagonal separates the two mutants, DomDIM and ΔSH3, from the other mutants as the first axis of the PCA for the SKAP2 data. The second diagonal separates the other mutants: a group of mutants with the lowest values in the two axes contains D120K, ΔLink2A, ΔLink2AB, W333K and that with the highest values Link2SH3, ΔLink1PH, ΔPH, ΔLink2B mutants. A similar organization was detected for the second axis of the PCA for the SKAP2 data.

Figure 3B shows that DomDIM and ΔSH3 have the highest value for FRK, YES, and FGR SRC kinases. Table 2 shows a summary of the data.

A hierarchical clustering followed by a consolidation using K mean method was performed on the two data taking account of the PCA to define the number of groups. Figure 5 shows the results. The hierarchical cluster tree and the factor map show that the three clusters are highly similar: cluster 3 of SKAP2 data and cluster 1 of SKAP1 data contain mutants in which the DIM domain is present and the SH3 domain deleted; both cluster 2 of SKAP2 and SKAP1 data contain D120K, ΔLink2A, ΔLink2AB, W330K/W336K mutants; and cluster 1 of SKAP2 data and cluster 3 of SKAP1 data. Link2SH3, ΔPH, ΔLink1PH, ΔLink2B, and DIM^-^ mutants. These very similar results strongly support that the two patterns detected in SKAP2 data exist also in SKAP1 even if the variability explains by the pattern defined by deletion of the SH3 domain and presence of the DIM domain is lower than in SKAP2. The two last clusters suggest a link between D120K/D129K and ΔLink2A and another one between DIM^-^ and ΔLink2B. An interesting hypothesis is that the two motifs A and B have completely different functions modulated by their phosphorylation status. The loss of intramolecular interaction between the DIM and the PH domains and functions associated with dimerization will be associated with an inactive motif A and the opposite is true for an inactive motif B.

In conclusion, these results support that for both proteins each module recognizes differently each kinase independently to other modules and that these patterns are globally conserved since their duplication. The main differences between these two proteins are quantitative: the diversity of the first pattern is stronger than that of the second one for SKAP2 in contrast to SKAP1 in which they have similar strength.

Figure 5: Factor map and hierarchical cluster tree from SKAP2 (A) and SKAP1 (B) data.

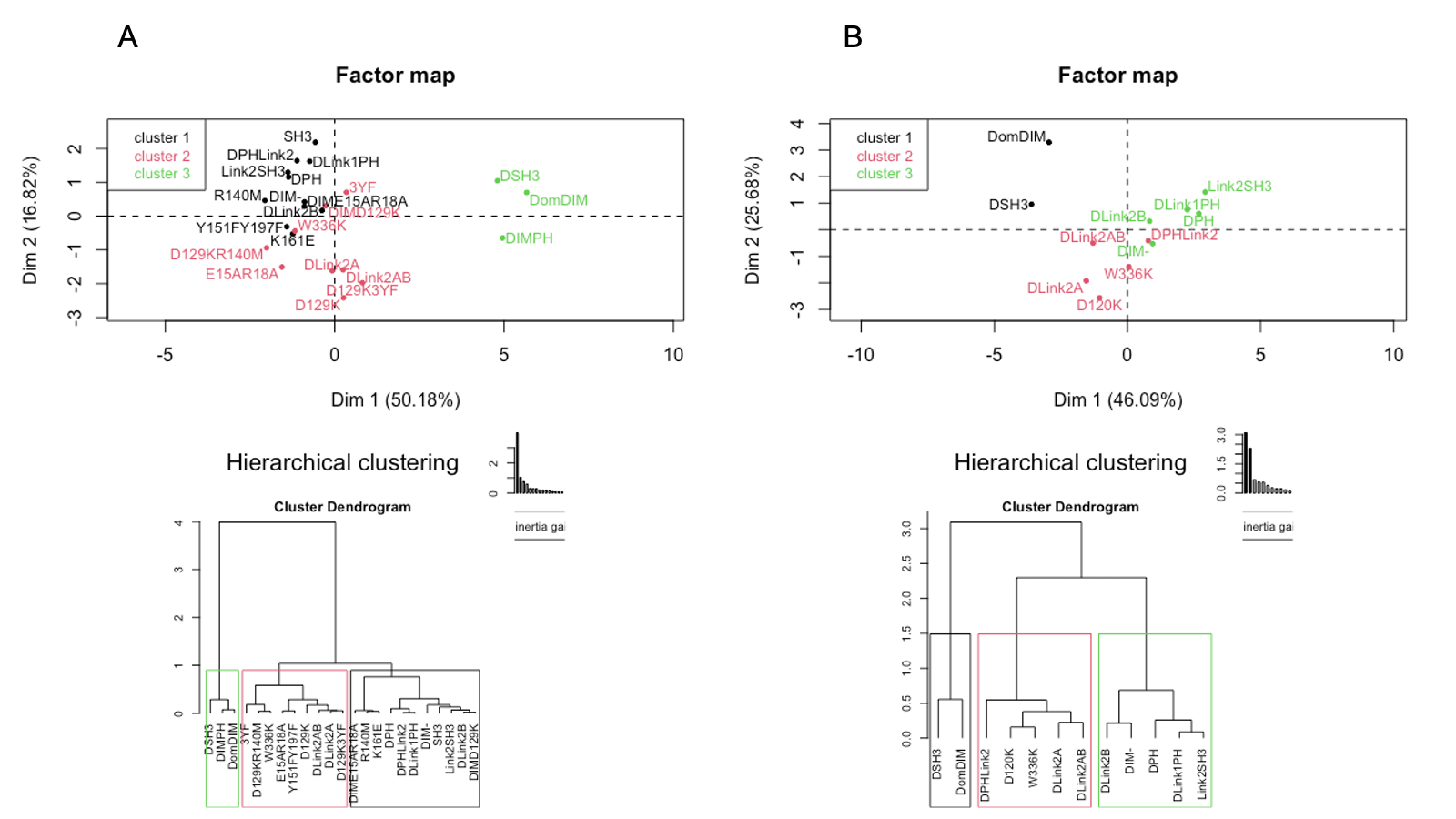

Table 1 Summary of the PCA for SKAP2 data

```
Eigenvalues
                       Dim.1   Dim.2   Dim.3   Dim.4   Dim.5   Dim.6   Dim.7   Dim.8   Dim.9
Variance               4.516   1.514   1.116   0.927   0.406   0.313   0.130   0.079   0.000
% of var.             50.178  16.823  12.396  10.300   4.507   3.474   1.445   0.877   0.000
Cumulative % of var.  50.178  67.001  79.397  89.697  94.204  97.678  99.123 100.000 100.000

Individuals (the 20 first)
                Dist    Dim.1    ctr   cos2    Dim.2    ctr   cos2    Dim.3    ctr   cos2
3YF         |  2.373 |  0.346  0.115  0.021 |  0.703  1.418  0.088 |  0.438  0.748  0.034 |
D129K       |  3.092 |  0.261  0.066  0.007 | -2.419 16.800  0.612 | -1.613 10.144  0.272 |
D129K3YF    |  2.433 |  0.819  0.646  0.113 | -1.974 11.191  0.658 | -0.388  0.587  0.025 |
D129KR140M  |  2.733 | -2.009  3.886  0.540 | -0.940  2.537  0.118 | -0.447  0.777  0.027 |
DIM         |  3.086 | -0.889  0.760  0.083 |  0.421  0.509  0.019 | -1.920 14.371  0.387 |
DIMD129K    |  1.369 | -0.266  0.068  0.038 |  0.303  0.264  0.049 | -0.822  2.633  0.361 |
DIME15AR18A |  2.865 | -0.894  0.770  0.097 |  0.286  0.235  0.010 |  2.023 15.951  0.499 |
DIMPH       |  5.162 |  4.955 23.638  0.921 | -0.647  1.203  0.016 |  0.564  1.240  0.012 |
DLink1PH    |  2.014 | -0.738  0.525  0.134 |  1.622  7.552  0.648 |  0.420  0.688  0.044 |
DomDIM      |  5.869 |  5.665 30.895  0.931 |  0.698  1.399  0.014 |  1.078  4.533  0.034 |
DPH         |  2.442 | -1.378  1.829  0.319 |  1.297  4.832  0.282 |  0.449  0.786  0.034 |
DPHLink2    |  2.251 | -1.112  1.190  0.244 |  1.639  7.715  0.530 |  0.437  0.743  0.038 |
DSH3        |  5.277 |  4.802 22.202  0.828 |  1.047  3.147  0.039 | -1.215  5.756  0.053 |
E15AR18A    |  2.637 | -1.557  2.333  0.348 | -1.508  6.529  0.327 | -0.595  1.378  0.051 |
K161E       |  2.425 | -1.236  1.471  0.260 | -0.527  0.797  0.047 |  1.704 11.310  0.494 |
Link2SH3    |  2.283 | -1.359  1.778  0.354 |  1.156  3.837  0.256 | -0.669  1.747  0.086 |
R140M       |  2.810 | -2.058  4.077  0.536 |  0.460  0.609  0.027 |  1.616 10.172  0.331 |
SH3         |  2.956 | -0.570  0.313  0.037 |  2.185 13.705  0.546 | -1.699 11.255  0.330 |
W336K       |  2.638 | -1.171  1.319  0.197 | -0.441  0.558  0.028 |  0.025  0.002  0.000 |
Y151FY197F  |  1.599 | -1.413  1.922  0.781 | -0.316  0.287  0.039 | -0.173  0.117  0.012 |

Variables
               Dim.1    ctr   cos2    Dim.2    ctr   cos2    Dim.3    ctr   cos2
HCK         |  0.657  9.554  0.431 |  0.509 17.129  0.259 | -0.118  1.255  0.014 |
LCK         | -0.793 13.915  0.628 | -0.350  8.101  0.123 |  0.203  3.691  0.041 |
BLK         | -0.741 12.157  0.549 |  0.507 17.006  0.257 |  0.207  3.858  0.043 |
LYN         |  0.217  1.038  0.047 | -0.543 19.440  0.294 |  0.237  5.039  0.056 |
FGR         |  0.341  2.579  0.116 |  0.141  1.312  0.020 |  0.875 68.633  0.766 |
YES         |  0.932 19.251  0.869 | -0.007  0.004  0.000 | -0.207  3.830  0.043 |
FYN         | -0.645  9.203  0.416 | -0.550 19.962  0.302 | -0.250  5.600  0.062 |
SRMS        | -0.786 13.683  0.618 |  0.488 15.697  0.238 | -0.263  6.209  0.069 |
FRK         |  0.917 18.620  0.841 | -0.143  1.349  0.020 | -0.145  1.885  0.021 |

Supplementary categories
                Dist    Dim.1   cos2 v.test    Dim.2   cos2 v.test    Dim.3   cos2 v.test
Del         |  1.731 |  1.283  0.550  2.068 |  1.124  0.422  3.130 | -0.079  0.002 -0.258 |
Exp         |  1.152 | -0.066  0.003 -0.057 | -1.015  0.776 -1.498 |  0.263  0.052  0.452 |
Mut         |  1.114 | -0.965  0.750 -1.555 | -0.453  0.166 -1.262 | -0.065  0.003 -0.211 |
Mut2        |  1.094 | -0.588  0.289 -0.595 | -0.581  0.282 -1.017 |  0.092  0.007  0.187 |
```

Table 2 Summary of SKAP1 data

```
Eigenvalues
                       Dim.1   Dim.2   Dim.3   Dim.4   Dim.5   Dim.6   Dim.7   Dim.8   Dim.9
Variance               4.148   2.311   1.060   0.582   0.398   0.333   0.138   0.030   0.000
% of var.             46.090  25.679  11.778   6.467   4.420   3.701   1.532   0.332   0.000
Cumulative % of var.  46.090  71.770  83.548  90.015  94.436  98.136  99.668 100.000 100.000

Individuals
             Dist    Dim.1    ctr   cos2    Dim.2    ctr   cos2    Dim.3    ctr   cos2
D120K    |  3.068 | -1.048  2.205  0.117 | -2.567 23.755  0.700 | -0.437  1.504  0.020 |
DIM-     |  2.404 |  0.942  1.783  0.154 | -0.527  1.001  0.048 |  1.811 25.786  0.568 |
DLink1PH |  2.654 |  2.246 10.130  0.716 |  0.748  2.017  0.079 |  0.233  0.428  0.008 |
DomDIM   |  4.547 | -2.947 17.450  0.420 |  3.292 39.068  0.524 | -1.017  8.128  0.050 |
DPH      |  3.361 |  2.681 14.437  0.636 |  0.608  1.334  0.033 |  0.452  1.609  0.018 |
DPHLink2 |  2.449 |  0.781  1.226  0.102 | -0.416  0.624  0.029 | -2.119 35.296  0.748 |
DSH3     |  4.159 | -3.605 26.102  0.751 |  0.954  3.281  0.053 |  1.693 22.523  0.166 |
Link2SH3 |  3.369 |  2.917 17.099  0.750 |  1.416  7.226  0.177 | -0.018  0.003  0.000 |
W336K    |  1.731 |  0.050  0.005  0.001 | -1.396  7.028  0.651 |  0.464  1.689  0.072 |
DLink2A  |  2.886 | -1.548  4.813  0.288 | -1.926 13.376  0.446 | -0.423  1.404  0.021 |
DLink2B  |  1.959 |  0.827  1.373  0.178 |  0.320  0.370  0.027 | -0.280  0.616  0.020 |
DLink2AB |  1.997 | -1.296  3.376  0.421 | -0.506  0.922  0.064 | -0.359  1.014  0.032 |

Variables
            Dim.1    ctr   cos2    Dim.2    ctr   cos2    Dim.3    ctr   cos2
HCK      |  0.568  7.766  0.322 |  0.695 20.906  0.483 | -0.122  1.414  0.015 |
LCK      |  0.191  0.881  0.037 | -0.824 29.361  0.679 | -0.035  0.114  0.001 |
BLK      |  0.829 16.556  0.687 |  0.235  2.388  0.055 | -0.331 10.322  0.109 |
LYN      |  0.279  1.876  0.078 |  0.152  0.994  0.023 |  0.941 83.534  0.886 |
FGR      | -0.737 13.088  0.543 |  0.457  9.021  0.208 | -0.097  0.879  0.009 |
YES      | -0.919 20.367  0.845 |  0.185  1.477  0.034 | -0.137  1.759  0.019 |
FYN      |  0.019  0.009  0.000 | -0.831 29.897  0.691 | -0.103  0.997  0.011 |
SRMS     |  0.904 19.715  0.818 |  0.288  3.581  0.083 | -0.100  0.950  0.010 |
FRK      | -0.905 19.742  0.819 |  0.234  2.375  0.055 |  0.018  0.030  0.000 |

Supplementary categories
             Dist    Dim.1   cos2 v.test    Dim.2   cos2 v.test    Dim.3   cos2 v.test
Del      |  1.212 |  0.346  0.081  0.563 |  1.100  0.824  2.400 | -0.129  0.011 -0.416 |
Exp      |  1.346 | -0.673  0.250 -0.632 | -0.704  0.273 -0.887 | -0.354  0.069 -0.658 |
Mut      |  1.710 | -0.019  0.000 -0.017 | -1.497  0.766 -1.885 |  0.612  0.128  1.139 |
```
